# HERC1 Loss of Function Causes a Developmental Disorder with Impaired Hippocampal Function

**DOI:** 10.64898/2026.10.30.763099

**Authors:** Eva M. Pérez-Villegas, Manel Garcia-Diez, Laura Costa-Sastre, Jose Manuel Hernández-Curiel, Guillem Verdaguer-Farré, Pau Sol Culleré, Francisco José López-Murcia, Angel M. Carrión, Francesc Ventura, José A. Armengol, Jose Luis Rosa

**Affiliations:** Department of Physiology, Anatomy and Cell Biology, Universidad Pablo de Olavide, 41013 Seville, Spain; Department of Physiological Sciences, University of Barcelona, IDIBELL, 08907 L’Hospitalet de Llobregat, Spain; Department of Pathology and Experimental Therapy, Institute of Neurosciences, University of Barcelona, and Bellvitge Biomedical Research Institute (IDIBELL), L’Hospitalet de Llobregat, 08907 Barcelona, Spain

**Author notes:** **Correspondence to:** Jose Luis Rosa Department of Physiological Sciences, University of Barcelona C/Feixa Llarga s/n, Planta 4ª (lab 4114), 08907 L’Hospitalet de Llobregat, Spain. These authors contributed equally to this work (Eva M. Pérez-Villegas, Manel Garcia-Diez, Laura Costa-Sastre).

**Keywords:** Ubiquitin ligase, loss of function, neurodevelopmental disorder, hippocampus, synaptic plasticity, cognitive impairment

## Abstract

Pathogenic variants in HERC1 are associated with a syndromic neurodevelopmental disorder characterized by macrocephaly, developmental delay, intellectual disability, and psychomotor dysfunction. Although genetic and molecular studies suggest a loss-of-function mechanism, definitive *in vivo* evidence that HERC1 deficiency is sufficient to cause neurodevelopmental abnormalities is lacking. Using a HERC1 knockout mouse model, we investigated the behavioural, neuronal, and synaptic consequences of complete HERC1 loss.

HERC1 deficiency caused motor coordination and cognitive deficits in both sexes, while male mutants additionally displayed hyperactivity and increased anxiety-like behaviour. Histological analyses revealed selective hippocampal abnormalities, including disruption of dentate gyrus organization, whereas Purkinje cells survival and cerebellar compartmentalization remained preserved. HERC1-deficient mice exhibited a marked increase in hippocampal GABAergic interneurons, impaired maturation of dendritic spines in CA1 pyramidal neurons, and altered adult neurogenesis, indicating widespread defects in hippocampal circuit organization.

Consistent with these structural abnormalities, glutamatergic synaptic components were significantly altered throughout the hippocampus. These molecular changes showed a sex-dependent pattern, with predominantly presynaptic defects in males and postsynaptic alterations in females. Despite these differences, whole-cell recordings revealed a common functional phenotype in both sexes. At both mossy fibre-CA3 and Schaffer collateral-CA1 synapses, HERC1-deficient neurons exhibited a progressive inability to sustain excitatory neurotransmission during repetitive stimulation, resulting in impaired steady-state glutamatergic transmission.

Together, these findings demonstrate that loss of HERC1 function is sufficient to produce a neurodevelopmental phenotype and identify HERC1 as a critical regulator of hippocampal circuit assembly and function. Our results provide mechanistic insight into the pathogenesis of HERC1-associated disorders and establish the *Herc1* knockout mouse as a preclinically relevant model for investigating disease mechanisms and therapeutic approaches.

## Introduction

Homozygous variants in the E3 ubiquitin ligase HERC1, including p.Ile469Aspfs*33,^1^ p.Leu1636Argfs*24,^2^ p.Ala2894Serfs*30,^3^ p.Arg3250*,^4^ and p.Arg4691Pro,^5^ have been associated with neurodevelopmental disorders.^6^ In addition, the co-occurrence of two heterozygous variants, p.Trp875* and p.Gly4520Glu, has also been linked to a neurodevelopmental phenotype.^7^ Comparison of these cases reveals a consistent clinical spectrum characterized by biallelic HERC1 deficiency, macrocephaly, dysmorphic facial features, developmental delay, intellectual disability, and impaired psychomotor development.^1,6^

To investigate HERC1 function *in vivo*, the *tambaleante* mouse model has been extensively characterized. This model carries a spontaneous missense mutation, p.Gly483Glu, within the RCC1-like domain of HERC1.^8^ The mutant protein accumulates at abnormally high levels which leads to reduced mTOR signalling and excessive autophagy. Phenotypically, *tambaleante* mice exhibit progressive Purkinje cell degeneration, severe cerebellar ataxia, reduced body size, impaired motor coordination, and shortened lifespan.^8^ Subsequent studies further demonstrated hippocampal and amygdala’s synaptic dysfunction associated with deficits in associative learning and memory.^9,10^

Although the *tambaleante* model has provided important insights into HERC1 biology, its pathogenic mechanism differs substantially from that underlying human disease. Most disease-causing HERC1 variants identified in patients are predicted to impair protein function through nonsense mutations, frameshifts, or missense substitutions affecting highly conserved residues within the catalytic HECT domain.^1,6^ Moreover, a mutant protein has been reported to be unstable and prone to degradation.^4^ In contrast, the *tambaleante* mutation does not affect the HECT domain and instead promotes protein stabilization and accumulation,^8^ a mechanism that has not been described in individuals with HERC1-associated neurodevelopmental disorders.

Consequently, it remains unclear whether loss of HERC1 function is, by itself, sufficient to cause a neurodevelopmental disorder. Addressing this question requires an *in vivo* loss-of-function model that more faithfully recapitulates the genetic and molecular mechanisms observed in affected individuals. Such a model would not only establish the causal relationship between HERC1 deficiency and disease pathogenesis but also provide a powerful platform for defining the physiological functions of HERC1 in the nervous system. Furthermore, it would constitute a clinically relevant preclinical model for investigating the cellular and molecular basis of HERC1-associated neurodevelopmental disorders and for evaluating potential therapeutic strategies.

## Materials and methods

### Animal model

The *Herc1KO* mouse strain (C57BL/6N-*Herc1^em3(IMPC)Wtsi^/Wtsi*) used in this study has been previously described.^11^ Briefly, a Crispr/Cas9-mediated deletion resulted in a 191 bp deletion that included exon 8 of the *Herc1* gene located in chromosome 9. Mice had *ad libitum* access to food and water in an environment with a 12-h light/dark cycle set at 22°C–24°C. The Ethics Committee for Animal Experimentation of the University of Barcelona and the Generalitat of Catalunya approved all procedures.

### Behavioral analysis

A total of 20 control (10 males and 10 females) and 20 *Herc1KO* (10 males and 10 females) mice were used for these analyses.

### Locomotor activity and anxiety

To evaluate locomotor and exploratory activities, mice were placed for 15 minutes in an open field (38×21×15 cm) (Cybertec S.A.). This apparatus consisted of a walled platform containing infrared emitters and sensors (IR) coupled to an altimeter. The movement sensor was connected to a computer that recorded the number of times the mice interrupted the IR beams/min.^12^

As anxiety-like behavior,^13^ the thigmotaxis; that is: the time that mice remained close to the wall of the open field without exploring the arena, was evaluated.

### Motor coordination

In addition to the easy protocol to test walk balance proposed by Guyenet et al.^14^ in which hind limb clasping, and gait were measured; two specific tests were used in present experiments.

### Static rods

Test was performed according to Deacon, ^15^ by using five 60 cm long wooden rods [35 (first rod), 28, 22, 15 and 9 (last rod) mm diameter each] fixed to the bench 60 cm above the floor. The finishing line was marked 10 cm on the extreme of the rod near the bench.

Mice were placed at the free end of the rod (nose tip oriented to the rod’s end). Time taken by mice to orientate from the starting position towards the bench (orientation time), and the time that mice employed to travel till finishing line (transit time) were measured. Test begun from the wide rod and finished by the smallest one. Both times were measured for each different size rods.

### Tail suspension test (TST)

TST was performed according to Can et al.^16^ procedure to analyze the tail-climbing behavior. Time in which mice climb their own tails, indicative that mice have learned that escape is possible, was measured. The maximum time for this test was 6 minutes.

### Nociception

The hot plate test used here to determine nociception was reported elsewhere. ^9^ In short, mice were placed on a hot plate maintained at 45, 50, and 55 °C (±0.7 °C). The pain behavioral response was measured as the time until the animal reared three times, leaning against the wall or jumping. The cutoff was 240 s.

### Learning and memory

#### Step-Through Passive Avoidance Test

The passive avoidance test was performed in a chamber [47 × 18 × 26 cm^3^, manufactured by Ugo Basile)] constructed with one light and other dark compartments.^9^ Briefly, mice were confined to the light compartment of the chamber. 30 s later, the door between the dark and light compartments was opened. Once the mice entered the dark compartment the door closed automatically and the mice received an electrical stimulus (0.5 mA, 5 s) delivered through the metal floor. In the retention tests, the latency to enter the dark compartment (escape latency) is a measure of information learning or memory retention.

### Morris water maze test

The spatial learning and memory test was previously reported.^10^ A 100 cm width and 45 cm height circular pool was filled 13 cm depth with water (23 ± 2 °C) opaqued by the addition of 0.01% TiO_2_. Four arbitrary quadrants N, S, E and W points divided the pool, and a 7 cm of diameter platform was hidden 1.5 cm below the water surface in the NO quadrant. Each mouse realized 4 trials per day with an inter-trial interval of 30 min. In each trial (one trial per quadrant), the mice were placed in each quadrant with their nose facing the pool wall, and all experimental sessions were recorded with a digital camera. Time spent to reach the platform (escape latency) was the main variable used, considering the maximum trial time as 90 sec. In the training session (1^st^ day) mice were first placed in the NE quadrant and, in case that they do not reach successfully the platform they were manually guided to it. In the acquisition period (2^nd^ to 5^th^ days) it was considered that mice have correctly found the platform when remained at least 10 sec on board. The retention interval was of seven days. Thus, day 12^th^ from the beginning of the test’s mice were subject to a trial similar to previous ones, but in which the platform was omitted. The time spent by mice to attain the exact pool’s area in which the platform should be located was measured.

### T-maze test

The exploratory memory was tested in a T-maze for three consecutive days (10 trial per day and mouse). Trials for spontaneous alternations measurement were performed.^10^ Briefly, the mice were confined in the arm they choose first for 30 sec. Thereafter, all the doors and central partition of T-maze were removed, and the animals were left to choose freely between the two arms.

### Immunohistochemical procedures and image quantification

Twelve controls and the same number of *Herc1KO* mice for each sex were used for these analyses. Mice were deeply anesthetized with an overdose of pentobarbital (80 mg/kg i.p.) and perfused intracardially with 4% paraformaldehyde in PB. After dissection, the brains were fixed overnight in the same fixative at 4 °C and immersed in 30 % sucrose in PB at 4 °C until they sank. Serial coronal sections (30 µm thick) were obtained on a Leitz freezing microtome, collected in PBS, and immunostained using our routine procedure.^9^ The primary antibodies used were: a rabbit polyclonal antiserum against calbindin (1:10000, Swant, CB-38); a mouse monoclonal antiserum against zebrin II (1:100, a gift from R. Hawkes);^17^ a rabbit polyclonal antibody against GABA (1:250, GeneTex, GTX125988); a goat polyclonal antibody against doublecortin (1:250, Santa Cruz, sc-8066); a rabbit polyclonal antiserum against the presynaptic vesicle protein SV2A (1:500, Synaptic Systems, 119002); a mouse monoclonal antibody against the vesicular glutamate transporter 1 (VGLUT1, 1:500, Synaptic Systems, 135011); a rabbit polyclonal antibody against the GluA1 subunit of AMPA glutamate receptors (1:500, Synaptic Systems, 182003); and a mouse monoclonal antibody against GluN1 subtype of NMDA glutamate receptors (1:500, Synaptic Systems, 114011). The secondary antibodies used were Alexa Fluor® 594 donkey-anti-goat (1:500, Invitrogen A150132); Alexa Fluor® 594 donkey-anti-mouse (1:500, Invitrogen A21203); and Alexa Fluor® 488 donkey-anti-rabbit (1:500, Invitrogen A21206). The sections were counterstained with DAPI (1:5000, D9542, Sigma) and images were acquired on a Leica SP5 confocal laser scanning microscope or on an upright Zeiss Axioimager M1 microscope. Confocal laser images for each pair of tested antibodies were obtained in the same session and under the same parameters (Laser 405: Intensity 31% Gain 10%; Laser 488: Intensity 60% Gain 82%; and Laser 561: Intensity 88% Gain 100%). The figure plates were prepared using the Photoshop 8.0.1 (Adobe ®) software without any additional correction of the microphotographs.

The immunoreactivities were quantified as indicated by Perez-Villegas et al.^9^ Briefly, an alternating sequence of laser pulses was used to activate the different fluorescent probes during image acquisition. Images from control and *Herc1*-KO mice were obtained under similar laser intensities and photomultiplier voltages, and in a same session for each pair of antibodies (e.g. Doublecortin-GABA, SV2-VGLUT1, and GluA1 and GluN1). Quantification of the fluorescent labelling density was performed offline with ImageJ and the size of the areas measured was determined automatically by defining outline masks based on brightness thresholds from maximal projected confocal images. The control and *Herc1*-KO images, whose values were expressed in arbitrary units, were captured from hippocampal areas similar in size (*p* = 0.98).

CaBP and GABA immunoreactive neurons were counted in similar in size hippocampal regions. The sections thickness, greater than the medium diameter of counted neurons, exclude the possibility of biassed counts. Therefore, CaBP and GABA immunoreactive neurons were counted without correction factors (for a rev. see ref.^18^). CaBP neurons counts were made on sections of the entire hippocampus at the same rostro-caudal levels, and their sizes ranged from 488861±132660 µm^2^ in controls to 492343±83854 µm^2^ in *Herc1 KO* animals (*p* = 0.958). As for CaBP counts, CA1 sections of the same rostro-caudal level were used for GABA+ neuron counts. The mean size of CA1 area was 63240±1636 µm^2^ in control animals and 65783±8357 µm^2^ in *KO* ones (*p* = 0.495).

### Dendritic spine analysis

Three controls and three *Herc1KO* mice for each sex were used for this experiment. The Golgi-Cox procedure used here was previously reported.^10^ Briefly, 4% paraformaldehyde in phosphate buffer 0.1M (PB, pH 7.2-7.4) fixed brains were scised in two sagittal halves. Each brain hemisphere block was immersed in the Golgi-Cox solution (1.78 % potassium dichromate, 1.78% mercuric chloride and 1.78% potassium chromate in distilled water). After 2 weeks in darkness at room temperature, the brain hemispheres were cryoprotected in 25% sucrose in Tris buffered saline 0.1M pH 7.4 (TBS) (24 hours at 4 °C). Sagittal sections (90 µm thick) were obtained on a Leitz freezing microtome and collected in TBS. Sections were treated with 0.1% Triton X-100 in TBS for 5 minutes, mounted on gelatine-coated slides and air-dried in the dark at room temperature. Dry sections were rinsed in distilled water (1 min) and immersed in 25% ammonium hydroxide solution (Fluka, cat. 17093-1L). After rinsing in distilled water (1 min) the sections were transferred for 20 min in Kodak Professional rapid fixer solution A. Thereafter the sections were rinsed with distilled water (1 min) and dehydrated with ascending grades of ethanol (70%, 90 % 1 min each, and 100% 2 times of 5 min each), cleared in xylene (2 x 10 min) and mounted with DPX. Images were acquired with a Zeiss Axioimager M1 microscope, and the figure plates were made using the Photoshop 8.0.1 (Adobe ®) software without any additional correction.

Dendritic spines were analysed as described previously.^10^ Z-stacks of Golgi-Cox stained and completely filled secondary dendrites from CA1 pyramidal neurons (optical section thickness = 0.5 µm) were obtained using a 100x oil-immersion objective with a numerical aperture of 1.74. The image series were converted to RGB using the FijiImageJ software (W. Rasband, National Institutes of Health, http://imagej-nih.gov/ij/) and the image sequences were then analysed with the Reconstruct software (http://synapses.clm.utexas.edu). Dendritic length, width and the length/width ratio was measured, and the spines were categorized according to the steps 2 and 3 of Risher et al. procedure.^19^ The data obtained were processed statistically in Excel Microsoft® (steps 4-6 of Risher et al.^19^).

### Electrophysiology

Acute horizontal slices of postnatal (P60-90) mice of either sex were used for electrophysiology. Animals were perfused with ice-cold low-Ca^2+^ and high-sucrose artificial CSF (aCSF) containing (in mM): 87 NaCl, 2.5 KCl, 25 NaHCO_3_, 1.25 NaH_2_PO_4_, 25 glucose, 0.5 CaCl_2_, 7 MgCl_2_, 75 sucrose (pH 7.4, bubbled with 95% O_2_, 5% CO_2_). After decapitation, brains were dissected and immersed in the same solution. Brains were glued onto the stage of a VT1000S vibratome (Leica), and 290 µm-thick horizontal slices containing either the CA3 region or the CA1 region of the hippocampus were cut. Slices were incubated for 40 min at 35°C in a chamber containing the same solution as the one used for dissection and slicing. Slices were kept in this solution at RT (21–24°C) for up to 5 h after recovery.

Whole-cell patch-clamp recordings were made either from granule cells, pyramidal neurons of the CA3 region, or pyramidal neurons of the CA1 region of the hippocampus at near physiological temperature using an EPC-10 amplifier controlled by Pulse or PatchMaster software (HEKA Elektronik). Sampling intervals and filter settings were 20 μs and 5 kHz, respectively. For voltage-clamp recordings, the membrane potential was set to −70 mV. Slices were placed into the recording chamber and allowed to wash for 10 min in normal aCSF before recordings commenced. Normal aCSF contained (in mM): 125 NaCl, 2.5 KCl, 25 NaHCO_3_, 1.25 NaH_2_PO_4_, 25 glucose, 2 CaCl_2_, 1 MgCl_2_ (pH 7.4, bubbled with 95% O_2_, 5% CO_2_). Neurons were visualized using near IR illumination through a 40× (Zeiss Achroplan, 0.8 NA Water immersion) objective. Perfused solution was warmed by a Single Inline Solution Heater SH-27B, controlled by a Single-Channel Temperature Controller TC324C (Multichannel Systems).

Patch pipettes were pulled from borosilicate glass capillaries with filament (Science Products) to have an open-tip resistance of 3–5 MΩ when filled with intracellular solution. The intracellular solution contained (in mM): 120 K-gluconate, 30 KCl, 10 EGTA, 2 MgCl_2_, 2 Na-ATP, 10 HEPES, 2 QX-314, 314 mOsm, pH 7.3. We did not correct for the liquid junction potential.

During voltage-clamp recordings, aCSF was supplemented with 25 μM bicuculline methiodide (HelloBio) to block GABA_A_R-mediated IPSCs. Mossy fibers and Schaffer collateral fibers were stimulated with a glass electrode filled with aCSF and placed in stratum lucidum or CA1 stratum radiatum, respectively, ≥80 μm away from the cell body of the patched postsynaptic cell. Transmitter release was evoked by applying brief electrical pulses (140 μs, 10–50 V). Stimulus intensity was adjusted to obtain peak EPSC amplitudes corresponding to single-axon stimulation, assuming that mean putative unitary EPSC amplitudes have a range of 40–500 pA.

All offline analysis of electrophysiological data was performed in Igor Pro (Wavemetrics). Voltage-clamp recordings were leak-corrected and low-pass filtered with a cut-off frequency fc=3 kHz using a 10-pole digital Bessel filter.

### Statistical analysis

Data were analyzed using GraphPad Prism 8, Microsoft Excel or Igor Pro and R (R Foundation for Statistical Computing), and are presented as mean ± standard error indicated. Statistical comparisons between KO and WT mice were performed using a two-tailed Student’s t-test. Differences were considered statistically significant at *p* < 0.05. Levels of statistical significance are indicated as follows: \**p* < 0.05, \*\**p* < 0.01, and \*\*\**p* < 0.001.

## Results

### Pathogenic HERC1 variants suggest a critical role for HERC1 in neurodevelopment

Human pathogenic HERC1 variants are predominantly predicted to impair or abolish HERC1 function, leading to the hypothesis that HERC1 deficiency underlies the associated neurodevelopmental disorders (Supplementary Fig. 1). To directly test this hypothesis, we generated a HERC1 loss-of-function mouse model and examined the effects of complete HERC1 deficiency *in vivo* (Supplementary Fig. 2).^11^

### HERC1 deficiency impairs motor coordination and cognition and produces male-specific behavioural alterations

To determine whether HERC1 deficiency recapitulates key features of HERC1-associated neurodevelopmental disorders in humans, we performed comprehensive behavioural and cognitive phenotyping of *Herc1* knockout (*KO*) mice. Locomotor activity, anxiety-like behaviour, motor coordination, nociception, and cognition were assessed using established behavioural paradigms (Table 1).^8–10,12^ HERC1 deficiency resulted in significant impairments in motor coordination and cognitive performance in both male and female KO mice compared with their wild-type (WT) littermates, whereas nociceptive responses remained unaffected. Notably, male KO mice displayed increased locomotor activity and anxiety-like behaviour, indicating a sex-dependent role for HERC1 in the regulation of these behavioural traits. These findings suggest that loss of HERC1 function produces behavioural abnormalities that partially mirror the neurological manifestations observed in individuals with HERC1-associated neurodevelopmental disorders.

**Table 1.** Behavioral analysis of *Herc1* knockout mice.

| Behavioral tests | Parameter measure | Test values: mean ± error (n) |  |  |  |  |  |
|---|---|---|---|---|---|---|---|
|  |  | WT males | KO males | *P* value | WT females | KO females | *P* value |
| **Open field** | Locomotion | 5916± 243 (8) | 8787± 474 (9) | <0,001 *** | 7961± 698 (9) | 6790± 429 (8) | 0,186 |
|  | Anxiety | 91,0±1,5 (8) | 94,6±0,8 (9) | 0,046 * | 94,4±0,9 (9) | 93,5±1,1 (8) | 0,584 |
| **Walk balance** | Motor coordination | 13,35± 0,72 (7) | 8,55± 0,55 (9) | <0,001 *** | 15,00± 0.01 (9) | 12,71± 0.95 (7) | 0,022 * |
| **Tail suspension** |  | 3,62± 1,47 (8) | 26,22± 3,42 (9) | <0,001 *** | 5,00± 0,95 (9) | 25,12± 4,61 (8) | <0,001 *** |
| **Hot plate** | Nociception | 10,00± 0,74 (8) | 11,88± 0,87 (9) | 0,125 | 22,54± 1,26 (9) | 19,48± 2,02 (8) | 0,209 |
| **Passive avoidance** | Cognition | 4,11± 0,87 (8) | 0,66± 0,12 (9) | <0,001 *** | 9,17± 2,49 (9) | 1,49± 0,66 (8) | 0,012 * |
| **Water Morris maze** |  | 23,6± 1,6 (8) | 41,4± 10,7 (9) | 0,015 * | 26,5± 2,5 (9) | 60,6± 11,9 (7) | 0,003 ** |
| **T-Maze** |  | 1,35± 0,28 (8) | 1,06± 0,27 (9) | 0,073 | 2,08± 0,28 (9) | 0,71± 0,28 (7) | 0,006 ** |

### HERC1 deficiency disrupts dentate gyrus architecture while preserving Purkinje cell organization

The cerebellum and hippocampus are key brain regions involved in motor coordination and learning, respectively, and structural alterations in their architecture have been linked to motor and cognitive dysfunction.^20,21^ We therefore investigated whether HERC1 deficiency affects the organization of these structures. To this end, we performed immunohistochemical analyses using calbindin-D28k (CaBP) to visualize Purkinje cells and hippocampal neuronal populations, and Zebrin II to evaluate Purkinje cell patterning and cerebellar compartmentalization. Since the observed effects were similar in both sexes, data from male and female mice are presented collectively.

Our analyses revealed that HERC1 deficiency neither affects Purkinje cell survival nor disrupts their characteristic monolayer organization (Supplementary Fig. 3). Preservation of Purkinje cell integrity in *Herc1 KO* mice is further supported by the absence of degenerative changes in their axons (Supplementary Fig. 3) and by the maintenance of a normal pattern of Purkinje cell projections to the deep cerebellar nuclei (Fig. 1A, B). This contrasts with other HERC1 mutation that is associated with Purkinje cell degeneration.^22^ In contrast, CaBP immunohistochemistry revealed clear abnormalities in the dentate gyrus of HERC1-deficient mice. Specifically, the granule cell layer contained multiple CaBP-immunonegative patches distributed throughout its entire extent (Fig. 1C, D, arrows in D; Supplementary Fig. 4, arrowheads in B, D and F). Moreover, CaBP-immunoreactive (CaBP-Ir) cell counts performed in the entire hippocampus whose area size were similar in both mice (*p* = 0.958), shown that this disruption of granule cell layer organization was accompanied by an increase in the number of CaBP-Ir cells throughout the hippocampus of HERC1-deficient mice (Supplementary Fig. 4G). Numerous CaBP-Ir cells were observed within the pyramidal cell layer, stratum oriens, and stratum lacunosum-moleculare (Supplementary Fig. 4A, C, E). HERC1 deficiency also appeared to affect calbindin expression in the cerebral cortex, which displayed a qualitatively greater number of CaBP-Ir cells across all cortical layers, with particularly noticeable differences in the granular layer (layer IV) (Fig. 1C, D). However, these cortical alterations will require further quantitative analysis to determine their extent and significance.

**Figure 1.**
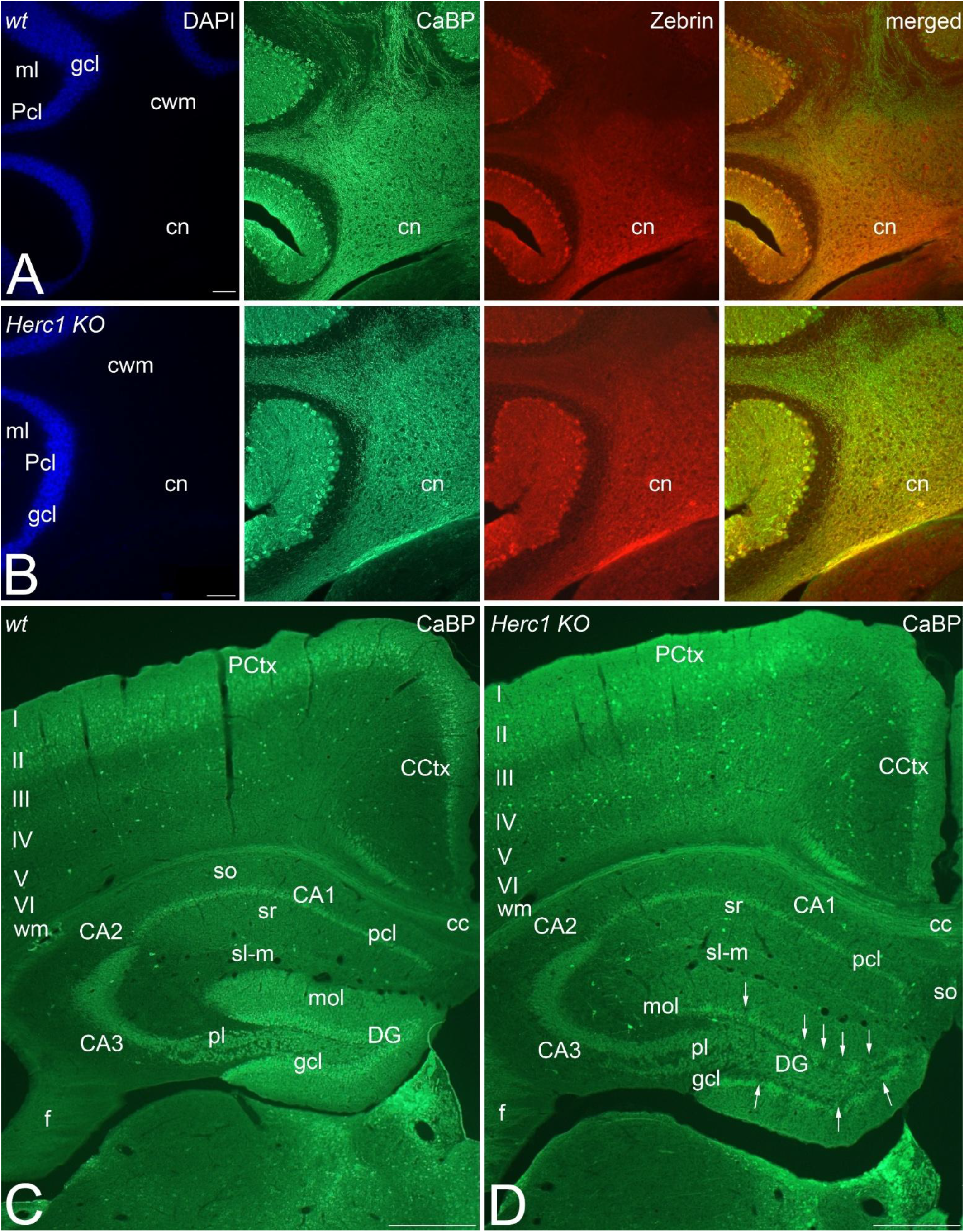
HERC1 regulates the structural organization of the dentate gyrus granule cell layer. Coronal sections through the brain (**C**-**D**) and the cerebellar nuclei (**A-B**, cn) of two-month-old mice. *Herc1 KO* does not affect the Purkinje cell axons coursing the cerebellar white matter (cwm) and their endings within the cerebellar nuclei (compare **A** to **B**). However, Calbindin (CaBP) immunoreactivity differences between control (**C**) and KO (**D**) cerebral cortex stand out. Layer IV of the *Herc1 KO* cerebral cortex qualitatively possesses more CaBP labelled neurons than that of control mice (compare **C** to **D**). In the dentate gyrus (DG) HERC1 deficiency disorganizes CaBP immunoreactivity of the granule cell layer (gcl) in which patches devoid of CaBP^+^ cells are evident (**D**, arrows), correlating with a fewer calbindin labelling of *KO* molecular layer (mol). CCtx, cingulate cerebral cortex; PCtx, parietal cerebral cortex; cc, corpus callosum. CA1, cornu ammonis 1; CA2, cornu ammonis 2; CA3, cornu ammonis 3; DG, dentate gyrus; S, subiculum; f, fimbria; pcl, pyramidal cell layer; pl, polymorphic cell layer; sl-m, stratum lucidum-moleculare; so, stratum oriens; sr, stratum radiatum; wm, white matter; I-VI, layers I to VI of the cerebral cortex. Bars = 1mm (**A**-**D**), and 200 µm (**C**-**D**). Similar results were observed in male and female mice (*p* > 0.05). Therefore, the data are presented collectively.

Collectively, these observations indicate that loss of HERC1 selectively compromises the organization of the hippocampus and, potentially, other cortical structures, while largely preserving the overall architecture of the cerebellum and the integrity of the Purkinje cell layer.

### HERC1 deficiency increases the number of GABAergic interneurons, limits adult neurogenesis, and alters dendritic spine maturation in hippocampal pyramidal neurons

GABAergic interneurons play a critical role in maintaining the excitatory-inhibitory balance within hippocampal circuits.^23,24^ We therefore investigated whether HERC1 deficiency affects hippocampal interneuron populations. As described above, *Herc1 KO* mice exhibited a marked and widespread increase in calbindin-immunoreactive (CaBP-Ir) neurons throughout the hippocampus (Supplementary Fig. 4D,F), raising the possibility that HERC1 deficiency may increase the number of interneurons. However, calbindin-positive interneurons represent only a subset of hippocampal inhibitory neurons and are not among the most abundant interneuron populations. Moreover, their functional roles relative to other interneuron subtypes remain incompletely understood.^25^ To determine whether this effect extended beyond calbindin-positive interneurons, we examined the broader population of inhibitory neurons by assessing GABA immunoreactivity throughout the hippocampus.^26^

*Herc1 KO* mice display a significant increase in the number of GABA-immunoreactive neurons in the hippocampus of both sexes compared with control animals (Fig. 2). GABA-positive neurons show the varied morphology described in their seminal works by Ramón y Cajal (see figs. 789 and 790 of ref.^27^), and are particularly abundant in the stratum oriens, and in the border between the stratum radiatum and the stratum lacunosum-moleculare in the hippocampus of *Herc1 KO* mice (Fig. 2B, arrows). In addition, GABA-immunoreactive somata consistent with basket cells,^28^ were readily detected within the pyramidal cell layer, superimposed on the characteristic diffuse GABA-immunoreactive background of this layer, whereas such somata were rarely observed in control animals (Fig. 2B, arrowheads). Quantitative analysis confirmed a statistically significant increase in the number of GABA-positive neurons in the CA1 region of *Herc1 KO* mice (Fig. 2C).

**Figure 2.**
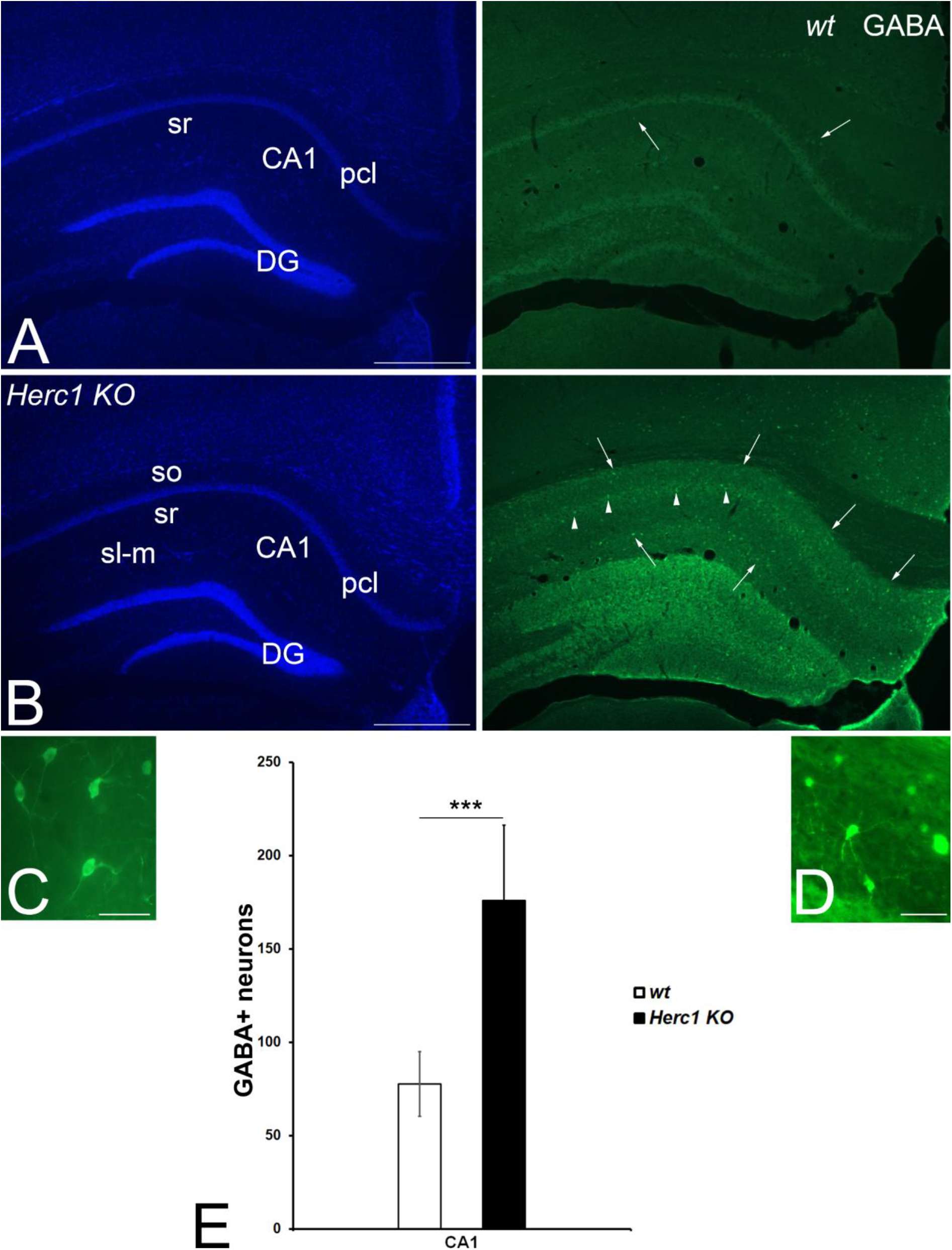
HERC1 deficiency increases the abundance of GABAergic neurons in the CA1 region of the hippocampus. Coronal sections through the brain of control **(A)** and *Herc1 KO* (**B**) mice. Gamma aminobutyric acid (GABA) immunoreactive neurons are most abundant in both CA1 hippocampal plexiform layers (so, stratum oriens; sr, stratum radiatum) of the *Herc1 KO* (**B**, arrows) than in the control hippocampus (**A**, arrows). In addition, GABA immunoreactive somata are found within the pyramidal cell layer (pcl) of *Herc1 KO* hippocampus (**B**, arrowheads), while lacking in control one (**A**). (**C)** and (**D)** illustrate the diversity in morphology of GABA interneurons (see also refs.^25,26^) counted here at the level of the stratum radiatum of CA1 region. Quantitative analysis demonstrates that the *Herc1 KO* CA1 has significantly more GABA immunoreactive neurons than the control (**E**; *p* = 0.000254). In present study any *p*-value less than 0.05 was considered significant, indicated as follows: \**p* < 0.05, \*\**p* < 0.01, and \*\*\**p* < 0.001. Bars = 500 µm (**A**-**B**), and 50 µm (**C**-**D**). Similar numbers of immunoreactive interneurons were obtained when compared male and female mice (*p* > 0.05). Therefore, the data are presented collectively.

To further evaluate whether the increased number of interneurons was accompanied by alterations in adult hippocampal neurogenesis, and to corroborate the behavioural findings, we analysed the immunohistochemical expression of doublecortin (DCX), a microtubule-associated protein expressed in immature neurons (Fig. 3). DCX immunoreactivity was assessed both as (i) a marker of adult hippocampal neurogenesis and (ii) a means of comparing the present findings with those previously reported in mice carrying the *tambaleante* HERC1 mutation.^10^ Since adult neurogenesis in the dentate gyrus gives rise predominantly to granule neurons,^29^ co-expression of GABA and DCX in neuronal somata would not be expected. However, confocal analysis revealed that a small subset of newly generated DCX-immunoreactive cells located in the subgranular zone adjacent to the granule cell layer also expressed GABA (Fig. 3). This observation is consistent with the findings of Liu et al., who reported the generation of GABAergic interneurons in the adult hippocampus.^30^ Although this finding lies beyond the scope of the present study, it provides additional support for the existence of adult-born GABAergic neurons. Notably, and in agreement with previous observations in *tambaleante* mice,^10^ we found a marked and statistically significant reduction in DCX immunoreactivity in the dentate gyrus of *Herc1 KO* mice compared with control animals (Fig. 3E), indicating impaired adult hippocampal neurogenesis associated with HERC1 deficiency.

**Figure 3.**
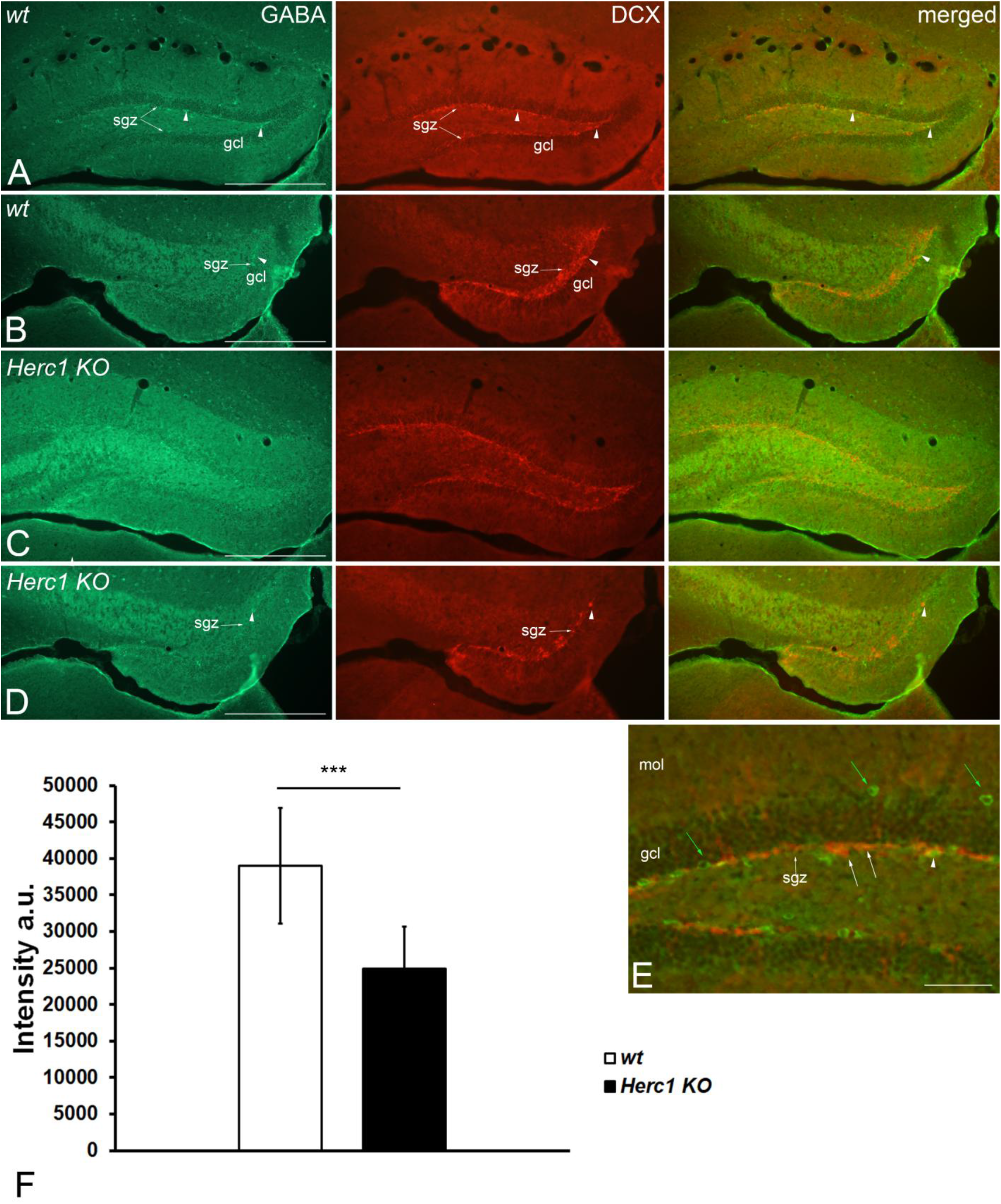
HERC1 deficiency impairs adult neurogenesis in the dentate gyrus. Coronal sections through the hippocampus of three-month-old control (**A**-**B**) and *Herc1 KO* mice (**C**-**D**). The expression of the microtubule-associated protein doublecortin (DCX) is lower in the dentate gyrus of *Herc1 KO* than in control mice. The majority of immunoreactive neurons only express GABA (**E**, green arrows) or DCX (**E**, arrows). However, it is possible to find GABAergic interneurons co-expressing DCX (**A**, **B**, **D**, **E**, arrowheads) located within the subgranular neurogenetic zone (sgz, small arrows) of the dentate gyri of both mice. Quantitative analysis demonstrates a stronger DCX immunoreactivity in the control dentate gyrus in relation to the *Herc1 KO* one (**F**, *p* = 0.000026). In present study any *p*-value less than 0.05 was considered significant, indicated as follows: \**p* < 0.05, \*\**p* < 0.01, and \*\*\**p* < 0.001. gcl, granule cell layer; mol, molecular layer. Bars = 500 µm (**A-D**) and 100 µm (**E**). Similar immunoreactive intensities were obtained when compared male and female mice (*p* > 0.05). Therefore, the data are presented collectively.

The increase in hippocampal inhibitory neurons was accompanied by abnormalities in the maturation of dendritic spines on CA1 pyramidal neurons (Fig. 4). Specifically, *Herc1 KO* mice exhibited a higher proportion of immature long-thin dendritic spines together with a reduced proportion of mature mushroom, stubby, and branched spines (Fig. 4J), a pattern indicative of impaired or delayed dendritic spine maturation. No sex-dependent differences in dendritic spine morphology were detected (Supplementary Figs. 5 and 6).

**Figure 4.**
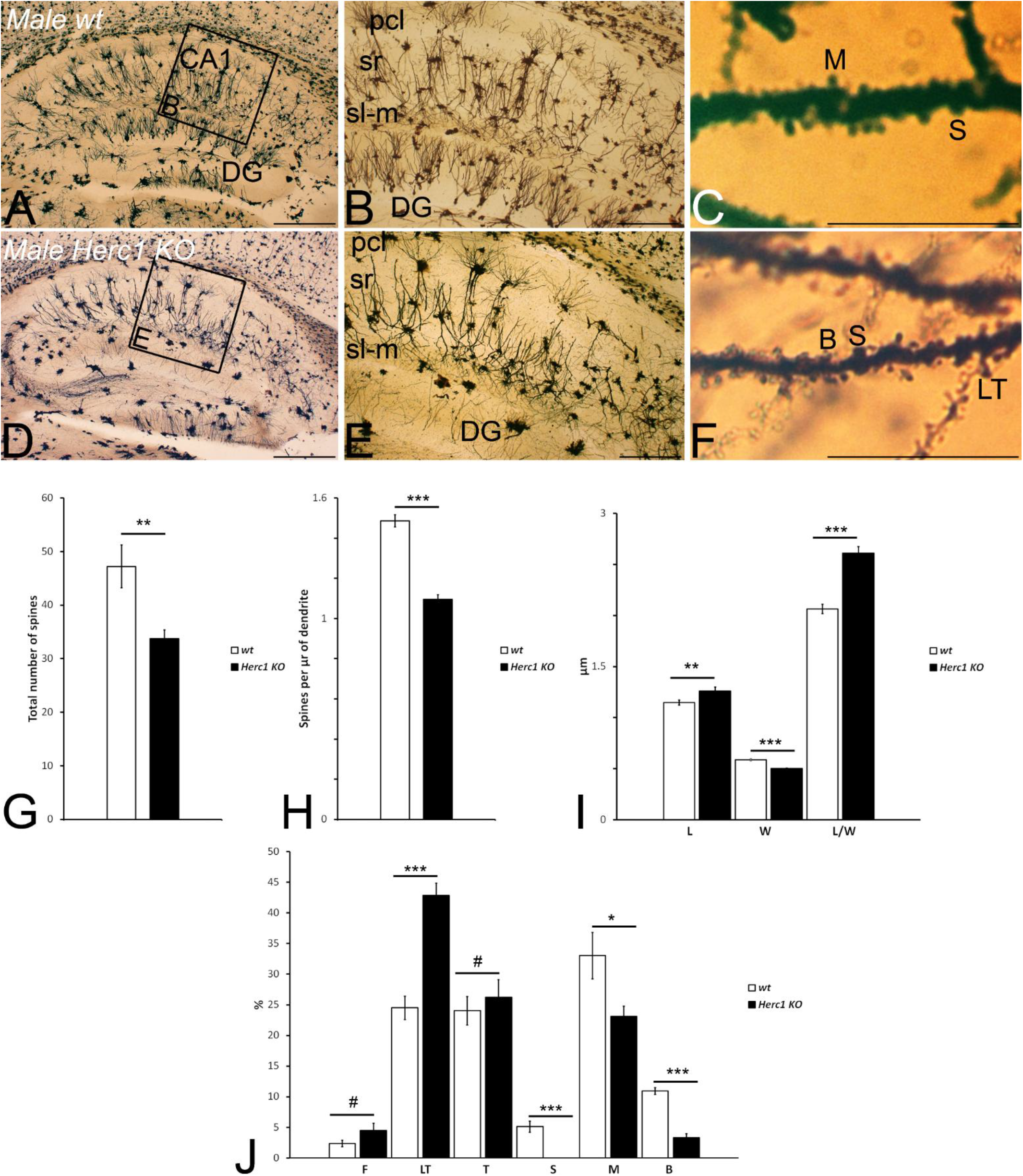
HERC1 regulates dendritic spine maturation in CA1 pyramidal neurons. Microphotographs of parasagittal sections through a Golgi-Cox impregnated hippocampus of control (**A-C**) and *Herc1 KO* (**D-F**) male mice. **C** and **F** illustrate the fully filled dendritic segments used to count and categorize the spines. **G to J**. Graphical representations of the number of spines counted (**G**, *p* = 0,005), the density of spines (**H**, *p* = 3,95^E-08^), and the spine differences in length (**I,** L, *p* = 0.034734), width (**I**, W, *p* = 9.3445^E-06^), and length/width ratio (**I**, L/W Ratio, *p* = 4.85531^E-06^) in the CA1 of wild-type and *KO* mice. The higher length/wide ratio indicates the mainly immature spines in the *KO* CA1. The proportions of the dendritic spine types counted are also shown in **J**. A significant decrease in the number of mushroom (**J**, M, *p* = 0.01269226), stubby (**J**, S, *p* = 0.00012968) and branched (**J**, B, *p* = 6.3083^E-07^) spines is observed in the *KO* mice (**J**). In addition, there is a significant increase in the number of immature long thin spines (**J**, LT, *p* = 4.9962^E-06^) in the *KO* CA1. The asterisks indicate significant differences in the Student’s t-test: \**p* < 0.05, \*\**p* < 0.01, and \*\*\**p* < 0.001. No significant values were indicated as follows: #*p* > 0.05. CA1, cornu ammonis 1; DG, dentate gyrus; F, filopodia; LT, long thin spines; T, thin spines; M, mushroom spines; S, stubby spines; B, branched spines. Bars = 500 µm (**A-B, D-E**), and 50 µm (**C, F**).

Taken together, these findings demonstrate that HERC1 deficiency affects both inhibitory and excitatory components of hippocampal circuitry. In contrast, the cerebellum, which is the primary target of the *tambaleante* mutation, appears structurally preserved in *Herc1 KO* mice.^8,22^ The concomitant increase in GABAergic interneurons, the reduction in adult neurogenesis, and the persistence of immature dendritic spines point to altered hippocampal development and circuit maturation. These observations, which were consistent across sexes, suggest that HERC1 deficiency disrupts the coordinated establishment of hippocampal neural networks, ultimately affecting the maturation of the principal hippocampal circuits.

### HERC1 deficiency impairs sustained glutamatergic transmission at hippocampal synapses

Based on the morphological alterations induced by HERC1 deficiency in the hippocampus, we asked whether HERC1 contributes to the functional maintenance of glutamatergic communication within hippocampal circuits. We examined synaptic transmission at the two major glutamatergic pathways of the hippocampus: mossy fiber and Schaffer collateral synapses.

We first focused on mossy fiber synapses, which originate from dentate gyrus granule cells (GCs) and terminate on the proximal dendrites of CA3 pyramidal neurons (CA3-Pyr) (Fig. 5A). Because changes in presynaptic neuronal excitability could potentially contribute to alterations in mossy fiber transmission, we first examined the intrinsic firing properties of GCs using whole-cell current-clamp recordings. GCs were subjected to 500 ms current injections ranging from 0 to +450 pA in 50 pA increments (Supplementary Fig. 7A). *Herc1 KO* GCs tended to generate action potentials at lower current intensities, consistent with increased initial excitability. However, at higher current intensities, *Herc1 KO* GCs exhibited action potential failures and reduced firing stability compared with control cells (Supplementary Fig. 7B–D). As comparable results were observed in male and female mice, the data are presented collectively.

**Figure 5.**
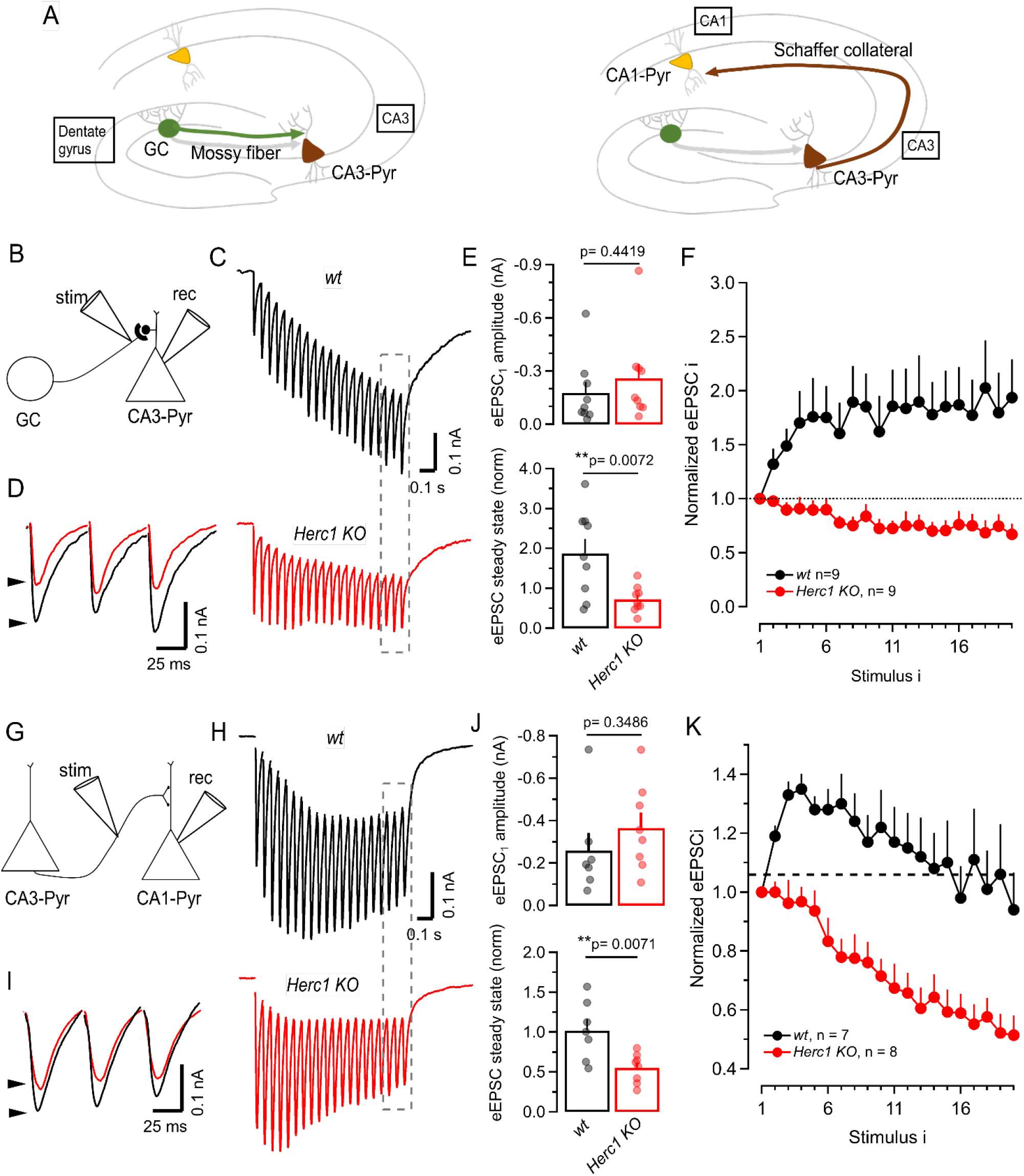
HERC1 deficiency impairs sustained glutamatergic transmission at hippocampal mossy fiber and Schaffer collateral synapses. (**A**) Schematic diagrams of hippocampal glutamatergic synaptic connections: (left) mossy fiber axons arising from dentate gyrus granule cells (GCs) and projecting onto the proximal dendrites of CA3 pyramidal neurons (CA3-Pyr); and (right) Schaffer collateral axons arising from CA3-Pyr neurons and projecting onto the proximal-to-distal dendrites of CA1 pyramidal neurons (CA1-Pyr). **(B**) Experimental electrophysiological configuration illustrating electrical stimulation of a putative unitary mossy fiber axon arising from a GC and whole-cell recording of postsynaptic responses from a CA3-Pyr neuron. **(C**) Mean evoked excitatory postsynaptic current (eEPSC) trains elicited by 20 stimuli at 20 Hz at hippocampal mossy fiber synapses in control (ctrl) and *Herc1 KO* mice. **(D**) Enlarged traces of the last three eEPSCs in the 20 Hz trains shown in (**C)**, corresponding to the regions indicated by striped rectangles. **(E**) Bar graphs and individual data points showing the amplitude of the first eEPSC (top) and the normalized steady-state eEPSC amplitude (bottom) during 20-stimulus trains at 20 Hz at hippocampal mossy fiber synapses in ctrl and *Herc1 KO* mice. For each synapse, steady-state eEPSC amplitude was calculated as the mean amplitude of the last five eEPSCs of the train and normalized to the amplitude of the first eEPSC. **(F**) Short-term plasticity at individual mossy fiber synapses, expressed as the amplitude of each eEPSC (eEPSC i) during 20 stimulus (stimulus i) trains at 20 Hz, normalized to the amplitude of the first eEPSC of the train. **(G**) Experimental electrophysiological configuration illustrating electrical stimulation of a putative unitary Schaffer collateral axon arising from a CA3-Pyr neuron and whole-cell recording of postsynaptic responses from a CA1-Pyr neuron. **(H-K**), Same as in (**C-F**), respectively, but for Schaffer collateral synapses. Error bars represent SEM (Student’s t-test, \*\**p*<0.01). As comparable results were observed in male and female mice, the data are presented collectively.

We next examined mossy fiber synaptic transmission by recording eEPSCs from CA3-Pyr neurons in response to stimulation of putative unitary mossy fiber axons (Fig. 5B). The amplitude of the first eEPSC was slightly larger in *Herc1 KO* than in control synapses, although this difference did not reach statistical significance (Fig. 5C, E *top*). In contrast, synaptic responses progressively diverged during the 20 stimulus, 20 Hz train, with *Herc1 KO* synapses exhibiting increasingly smaller eEPSCs than controls (Fig. 5C, D, F). Accordingly, the normalized steady-state response, calculated as the mean amplitude of the last five eEPSCs relative to the first eEPSC, was significantly reduced in *Herc1 KO* synapses (Fig. 5E *bottom*). Since the observed effects were similar in both sexes, data from male and female mice were pooled and are presented together.

We next investigated whether this phenotype also affected other major glutamatergic pathway of the hippocampal circuit, namely the Schaffer collateral-CA1 synapses (Fig. 5A,G). These synapses exhibited a pattern similar to that observed in the mossy fiber pathway. Although the amplitude of the first evoked EPSC tended to be larger in *Herc1 KO* than in control synapses, this difference did not reach statistical significance (Fig. 5H, J, top). In contrast, during stimulation at 20 Hz, synaptic responses progressively declined relative to controls, resulting in a significantly smaller steady-state response in *Herc1 KO* synapses (Fig. 5H-K).

These findings demonstrate that HERC1 deficiency progressively hampers the capacity of hippocampal glutamatergic synapses to sustain neurotransmission during repetitive activity, affecting both mossy fiber and Schaffer collateral pathways.

### HERC1 deficiency disrupts glutamatergic synapses in a sex-dependent manner

The structural and functional alterations observed in the hippocampus of *Herc1 KO* mice prompted us to investigate the molecular mechanisms underlying the impaired synaptic transmission. Glutamatergic neurotransmission constitutes the major excitatory component of the hippocampal trisynaptic circuit. We therefore analysed excitatory synapses in the dentate gyrus, CA3, and CA1 regions. Presynaptic terminals were examined using antibodies against synaptic vesicle protein 2A (SV2A), a general marker of synaptic vesicles,^31^ and vesicular glutamate transporter 1 (VGLUT1), a specific marker of glutamatergic presynaptic terminals.^32^ Postsynaptic components were assessed using antibodies against the GluA1 subunit of AMPA receptor,^33^ and the GluN1 subunit of NMDA receptors,^34^ two major mediators of excitatory synaptic transmission. Quantitative immunofluorescence analyses revealed region- and sex-specific alterations in the expression of glutamatergic synaptic markers.

In both male and female *Herc1 KO* mice, the presynaptic vesicle markers SV2A and VGLUT1 were significantly reduced in the dentate gyrus (Fig. 6), with the decrease being more pronounced in females (Fig. 6G). In male *Herc1 KO* mice, VGLUT1 expression was also reduced in the CA1 and CA3 regions compared with control animals, whereas SV2A immunoreactivity remained unchanged in the CA3 region (Supplementary Fig. 8). In contrast, female *Herc1 KO* mice exhibited a distinct pattern of presynaptic marker expression: SV2A levels were reduced in CA3 but remained unaltered in CA1, while VGLUT1 expression was preserved in CA3 and reduced in CA1 (Supplementary Fig. 9). Direct comparison between male and female *Herc1 KO* mice revealed generally lower levels of both presynaptic markers in females across all hippocampal regions analysed. However, the only statistically significant sex-dependent differences were observed for SV2A expression in the dentate gyrus (Fig. 6 **G**) and in the CA1 region (Fig. 7A), which was significantly lower in females than in males.

**Figure 6.**
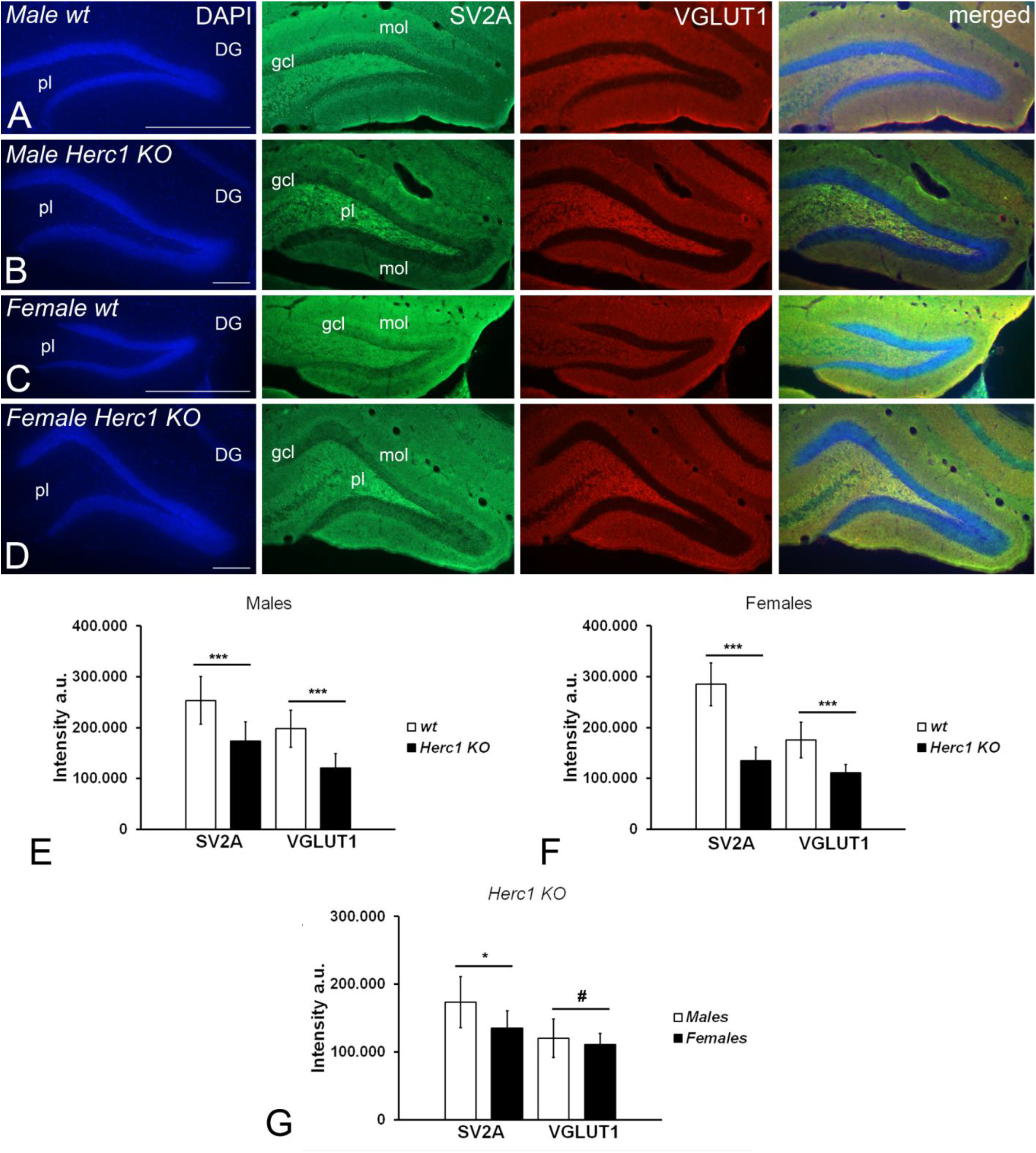
HERC1 deficiency disrupts presynaptic excitatory synapses. Coronal sections through the hippocampus of two-month-old male (**A**), female (**B**) control, and male (**C**) and female (**D**) *Herc1 KO* mice. Presynaptic proteins markers of glutamatergic synapses, the synaptic vesicle 2A (SV2A) and the vesicular glutamate transporter1 (VGLUT1), have a stronger immunoreactivity in the dentate gyrus of the control mice (**A**-**B**) compared to *KO* animals (**C**-**D**). Male *vs.* female qualitative differences in the immunoreactivity of both presynaptic proteins are evident at the pleomorphic layer of *KO* dentate gyrus (compare **B** and **D**). Quantitative analysis demonstrates the most intense SV2A and VGLUT1 immunoreactivity in the control dentate gyrus related to the *Herc1 KO* one (**E**, *p* = 0,0002 and *p* = 2,56^E-05^; **F**, *p* = 1,54^E-08^ and *p* = 7,84^E-05^). Moreover, there are sex-differences in the intensity of the immunoreactivity of both presynaptic proteins; thus, female *KO* dentate gyrus seems to possess less SV2A presynaptic protein (**G**, *p* = 0.0116) while the expression of presynaptic glutamate transporter VGLUT1 is under the level of significance (**G**, *p* = 0.3741). In present study any *p*-value less than 0.05 was considered significant, indicated as follows: \**p* < 0.05, \*\**p* < 0.01, and \*\*\**p* < 0.001. # *p*>0.05. DG, dentate gyrus; gcl, granule cell layer; mol, molecular layer; pl, polymorphic cell layer. Bars = 500 µm (**A**, **C**), and 200 µm (**B**, **D**).

**Figure 7.**
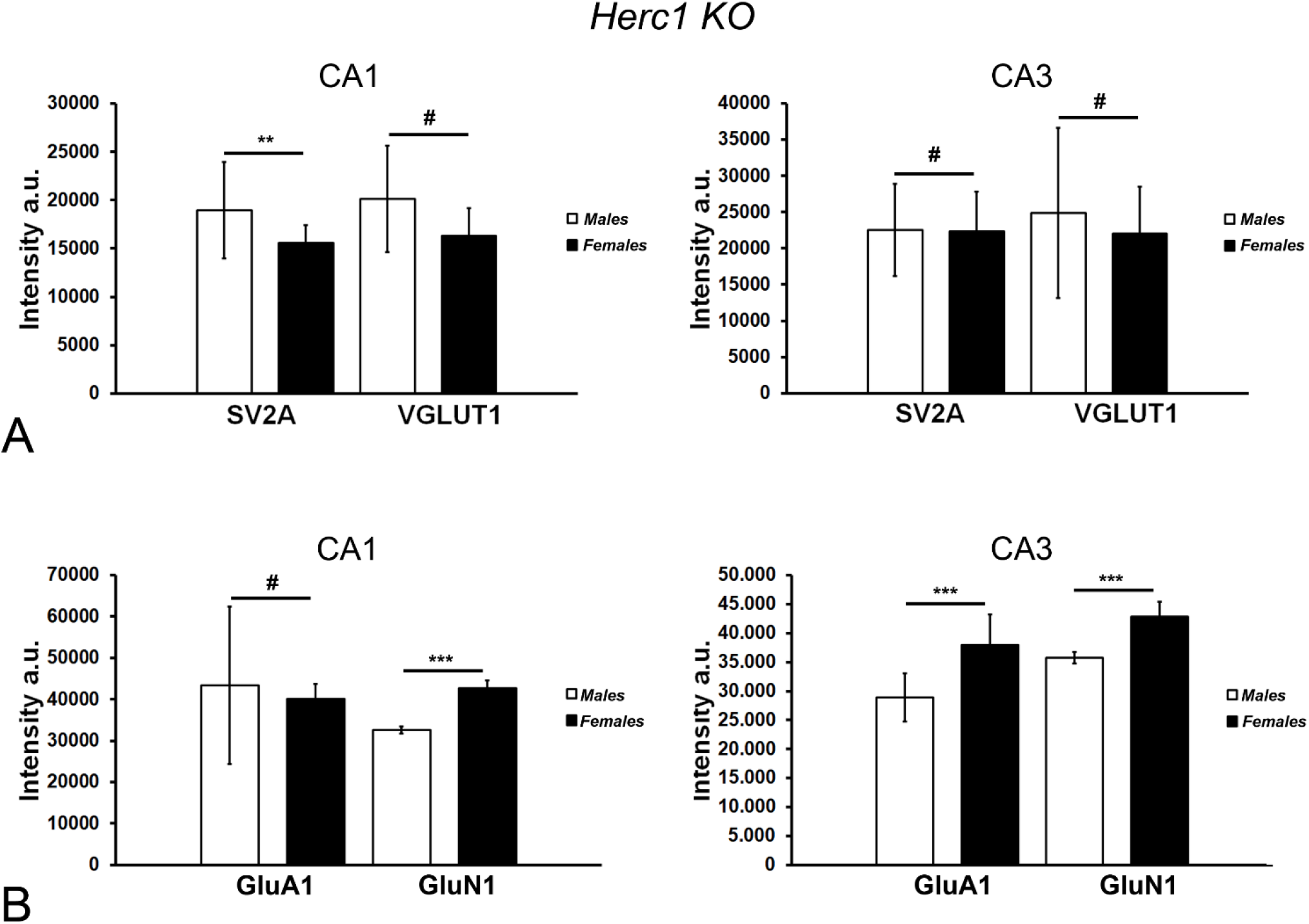
HERC1 deficiency induces sex-dependent alterations in pre- and postsynaptic proteins involved in excitatory neurotransmission. Graphics illustrate the differences in the expression of presynaptic **(A)** and postsynaptic **(B)** proteins in the CA1 and CA3 hippocampal areas. The only sex-difference of the synaptic vesicle protein 2 (SV2A) expression is observed in CA1 whose immunoreactivity is stronger, as it occurs in the dentate gyrus (see Fig. 7G), in male than in female *Herc1 KO* hippocampus (**A**, *p* = 0.003055). In contrast no significant sex-differences are found in VGLUT1 immunoreactivity intensities between CA1 (**A**, *p* = 0.8732) and CA3 (**A**, *p* = 0.8732) areas. Regarding the postsynaptic proteins, there are sex-differences in the expression of the subunit GluA1 of AMPA1 ionotropic glutamate receptor which is stronger in female CA3 than in male one (**B**, *p* = 1.02085^E-17^), remaining unaltered in CA1 (**B**, *p* = 0.3883). The expression of GluN1 subtype glutamate receptor was most intense in female CA1 (B, *p* = 1.02085^E-17^) and CA3 (**B**, *p* = 1.02058^E-16^) than in the same areas of male *Herc1 KO* hippocampus. The asterisks indicate significant differences in the Student’s t-test: \**p* < 0.05, \*\**p* < 0.01, and \*\*\**p* < 0.001. No significant values were indicated as follows: #*p* > 0.05.

We next examined the postsynaptic glutamatergic receptors GluA1 and GluN1. Female *Herc1 KO* mice exhibited a marked reduction in GluA1 expression throughout the dentate gyrus (Supplementary Fig. 10) as well as in the CA1 and CA3 regions (Supplementary Fig. 11). In contrast, GluN1 expression showed a region-specific pattern, being reduced in the dentate gyrus, increased in CA3, and unchanged in CA1 (Supplementary Figs. 10 and 11). In male *Herc1 KO* mice, the only significant alteration was an increase in GluN1 expression in the CA3 region (Supplementary Fig. 12), whereas no significant changes in either GluA1 or GluN1 levels were detected in the dentate gyrus (Supplementary Fig. 13) or CA1 region (Supplementary Fig. 12). Direct comparisons between male and female *Herc1 KO* mice revealed no significant sex-dependent differences in the expression of postsynaptic glutamate receptor markers in the dentate gyrus (Supplementary Fig. 14). However, in the CA1 and CA3 regions, female *Herc1 KO* mice exhibited significantly higher levels of postsynaptic glutamatergic markers than their male counterparts (Fig. 7B). These findings indicate that HERC1 deficiency differentially affects postsynaptic glutamatergic receptor expression according to both hippocampal subregion and sex.

Alterations in both presynaptic and postsynaptic glutamatergic markers, together with the increased number of calbindin-positive and GABA-immunoreactive interneurons, the reduction in adult neurogenesis within the dentate gyrus, and the persistence of immature dendritic spines in CA1 pyramidal neurons, demonstrate that HERC1 deficiency profoundly disrupts the structural and functional organization of hippocampal circuits. These abnormalities provide a cellular and molecular framework for the behavioural deficits observed in *Herc1 KO* mice. Although the molecular alterations affecting glutamatergic synaptic components differ between males and females, both sexes converge on a common neurobiological phenotype characterized by impaired excitatory connectivity and dysregulation of inhibitory neuronal networks. Together, these changes are likely to compromise hippocampal information processing and synaptic transmission, ultimately contributing to the cognitive and behavioural impairments associated with HERC1 deficiency.

## Discussion

The data presented in this study provide the first *in vivo* evidence that loss of function of the E3 ubiquitin ligase HERC1 causes cognitive and motor coordination deficits. Together with the macrocephaly phenotype previously reported in *Herc1 KO* mice,^35^ these findings establish this model as a valuable preclinical tool for investigating HERC1-associated neurodevelopmental disorders. Importantly, *Herc1 KO* mice recapitulate several key features observed in affected individuals, including neurodevelopmental abnormalities and impaired neurological function (Supplementary Fig.1).^1,6^ Beyond the behavioural phenotype, our results reveal profound alterations in hippocampal organization at multiple levels. HERC1 deficiency increased the numeric abundance of GABAergic interneurons, altered dendritic spine maturation, reduced adult neurogenesis, disrupted glutamatergic synaptic components, and impaired hippocampal neurotransmission. These abnormalities support a critical role for HERC1 in the establishment and maintenance of hippocampal circuitry.

Adult hippocampal neurogenesis has been directly linked to learning and memory processes.^29^ The marked reduction in doublecortin expression observed here highlights the detrimental effect of HERC1 deficiency on adult neurogenesis, consistent with previous findings in the *tambaleante* mouse model carrying a HERC1 mutation.^10^ Our behavioural analyses further show that, apart from preserved nociception and the absence of ataxia, the phenotype of *Herc1 KO* mice closely resembles that of *tambaleante* mice.^8–10,22^ Importantly, independently of the effects that altered glutamatergic transmission may have on learning and memory (see below), the profound impairment of adult hippocampal neurogenesis in *Herc1 KO* mice could itself contribute substantially to the behavioural deficits associated with HERC1 deficiency.^36^

An intriguing finding of the present study is the presence of sex-dependent alterations in glutamatergic synaptic markers despite the absence of sex-specific differences in dendritic spine morphology and in the number of calbindin and GABA expressing neurons. Male *Herc1 KO* mice predominantly exhibited presynaptic deficits, as indicated by reduced SV2A and VGLUT1 levels throughout the hippocampus, whereas female mutants displayed more pronounced postsynaptic alterations, characterized by decreased GluA1 and GluN1 expression. Glutamatergic synaptic transmission is widely recognized as a major contributor to the mechanisms underlying neural plasticity.^37^ Therefore, the increased abundance of GluA1 and GluN1 observed in the hippocampus of male *Herc1 KO* mice may reflect a compensatory plastic response of postsynaptic glutamatergic neurons to reduced synaptic input, similar to that described in other regions of the nervous system.^38^ In contrast, a stronger effect of HERC1 deficiency on glutamate receptor turnover in female mice could impair this compensatory mechanism, leading to the marked reduction in receptor expression observed in the female hippocampus. It is important to note that mossy fibers arising from dentate gyrus granule cells innervate GABAergic interneurons that regulate excitatory transmission within the CA3 region.^39^ Consequently, the increase in the number of GABAergic interneurons in the CA3 and polymorphic layers induced by HERC1 deficiency could alter hippocampal circuit activity and disrupt the normal synaptic output of CA3 pyramidal neurons through the Schaffer collateral pathway. Such alterations could impair hippocampal projections to the cingulate cortex,^25^ thereby contributing to the behavioural abnormalities observed in HERC1-deficient mice.^24^

Despite their molecular differences, both sexes exhibited comparable abnormalities in dendritic spine maturation, including an increased proportion of immature long-thin spines and a concomitant reduction in mature spine types. Furthermore, electrophysiological analyses revealed similar impairments in glutamatergic synaptic transmission in male and female KO mice. Together, these findings suggest that distinct sex-dependent molecular alterations converge on a common structural and functional phenotype, ultimately leading to comparable deficits in hippocampal synaptic connectivity and neurotransmission.

The basis of the sex-dependent molecular responses found here are unclear. Notably, circulating oestradiol levels were unchanged in both male and female *Herc1 KO* mice relative to their respective controls,^11^ arguing against a major contribution of altered oestrogen levels to the observed synaptic phenotype. In contrast, female *KO* mice exhibited significantly reduced testosterone and dihydrotestosterone levels, whereas no changes were detected in mutant males.^11^ Because androgens have been implicated in the regulation of hippocampal synaptic plasticity, glutamatergic neurotransmission, receptor trafficking, and dendritic spine maintenance,^40–44^ reduced androgen signalling may contribute to the differential expression of glutamatergic synaptic proteins observed in female mutants. Interestingly, sex-dependent effects of HERC1 deficiency were not restricted to molecular phenotypes. Male *KO* mice also exhibited increased locomotor activity and anxiety-like behaviours, suggesting that sex-specific mechanisms may influence the behavioural consequences of HERC1 loss of function. While the present work does not directly address the mechanisms underlying these behavioural alterations, future studies will be required to determine how the molecular and synaptic changes identified here ultimately affect neural circuit function and behaviour.

An important consequence of our findings is the demonstration that HERC1 plays a critical role in the regulation of glutamatergic neurotransmission. This observation raises the possibility that HERC1 dysfunction may contribute to neurological and neuropsychiatric disorders in which alterations of the glutamatergic system are implicated. Schizophrenia is a notable example, as multiple lines of evidence support a role for glutamatergic dysfunction, particularly NMDA receptor hypofunction, in disease pathogenesis.^45^ Consistent with this hypothesis, large-scale exome studies have identified ultra-rare coding variants in several glutamatergic genes, including GRIN2A and GRIA3, that substantially increase the risk of schizophrenia. Interestingly, HERC1 was also identified among these risk genes, and protein-truncating variants in HERC1 were associated with increased disease susceptibility.^46^ Moreover, the increase of GluN1 expression in female *Herc1 KO* CA3 correlates with that was observed in the brain of schizophrenic patients.^47^ Thus, despite that a conclusive relationship between HERC1 deficiency and schizophrenia remains to be established, our findings provide biological support for the hypothesis that disruption of HERC1-dependent glutamatergic signalling may contribute to the pathophysiology of neuropsychiatric disorders.

In summary, our results identify HERC1 as a key regulator of hippocampal circuit maturation, neuronal connectivity, and synaptic function. HERC1 deficiency induces a core neurodevelopmental phenotype characterized by altered interneuron populations, impaired dendritic spine maturation, reduced pyramidal neuron numbers, and defective synaptic transmission. While sex-dependent hormonal environments may modulate the molecular pathways through which these defects are expressed, resulting in distinct presynaptic and postsynaptic alterations, the overall structural and functional phenotype remains largely consistent between sexes. These findings provide mechanistic insight into the pathogenesis of HERC1-associated disorders and establish the *Herc1 KO* mouse as a valuable preclinical model for studying disease pathogenesis and developing therapeutic strategies for HERC1-associated neurodevelopmental disorders.

## Data availability

Any additional information reported in this paper is available from the lead contact upon request.

## Acknowledgements

We thank E. Adanero, A. Muhaisen, E. Castaño, B. Torrejon, the Biology-Bellvitge Unit from Scientific and Technological Centers (CCiTUB), Universitat de Barcelona for technical assistance. Skillful technical assistance of K. Garcia from the Centro Andaluz de Biología (CSIC-UPO) is also acknowledged. Manel Garcia-Diaz is a recipient of a STEP-FI fellowship from the Generalitat de Catalunya. Laura Costa-Sastre is a recipient of a PRE-DOCS UB fellowship from the Universitat de Barcelona.

## Supplementary material

Supplementary material is available at *Brain* online’.

---

**Supplementary Fig. 1.**
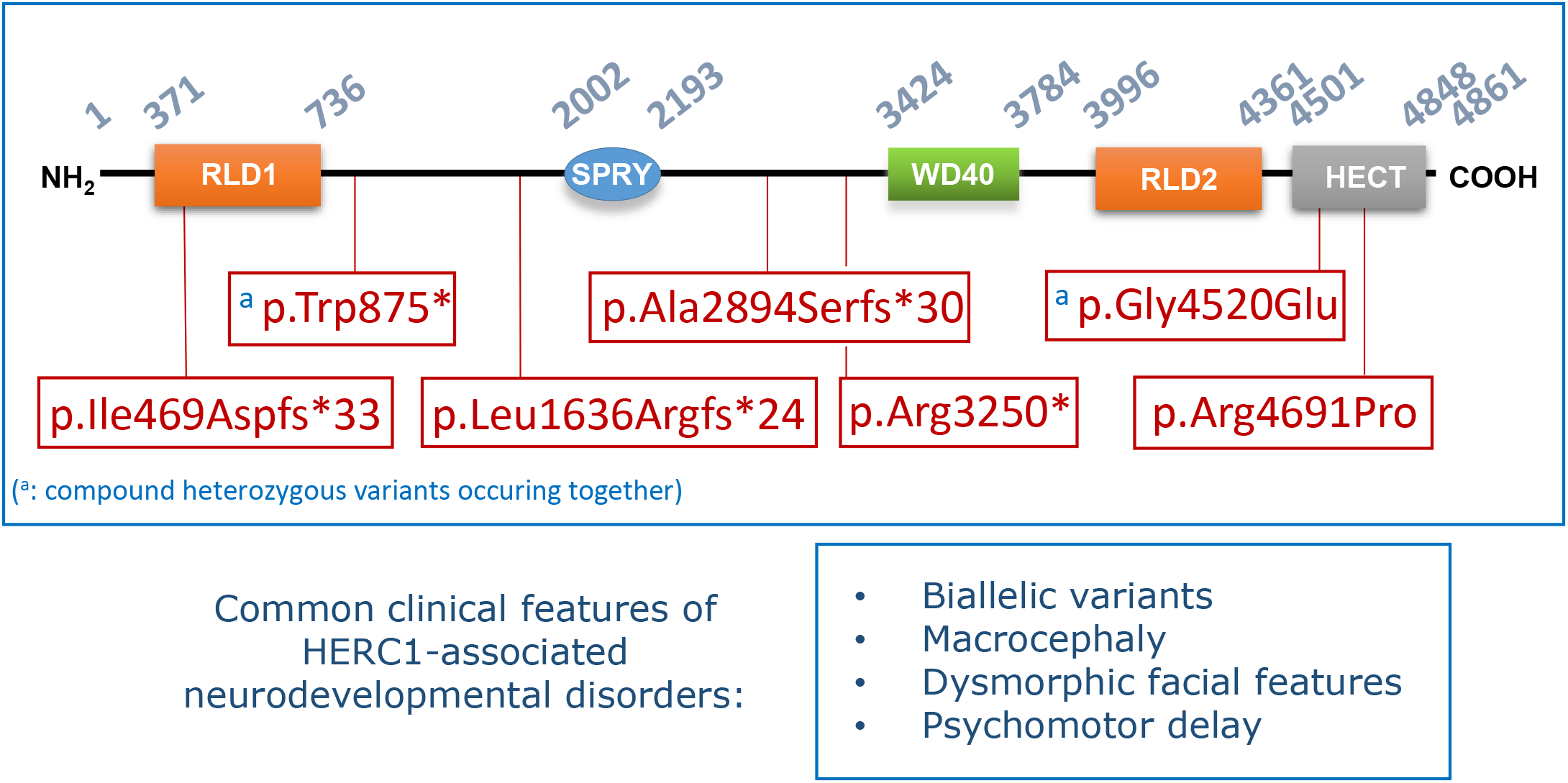
Pathogenic HERC1 variants associated with human neurodevelopmental disorders. Disease-associated HERC1 variants reported in individuals with neurodevelopmental disorders are mapped onto the HERC1 protein. Most variants are predicted to result in loss of function through protein truncation or disruption of key functional domains. Comparative analysis of these cases reveals a characteristic clinical spectrum marked by biallelic HERC1 deficiency, macrocephaly, dysmorphic facial features, and psychomotor delay.

**Supplementary Fig. 2.**
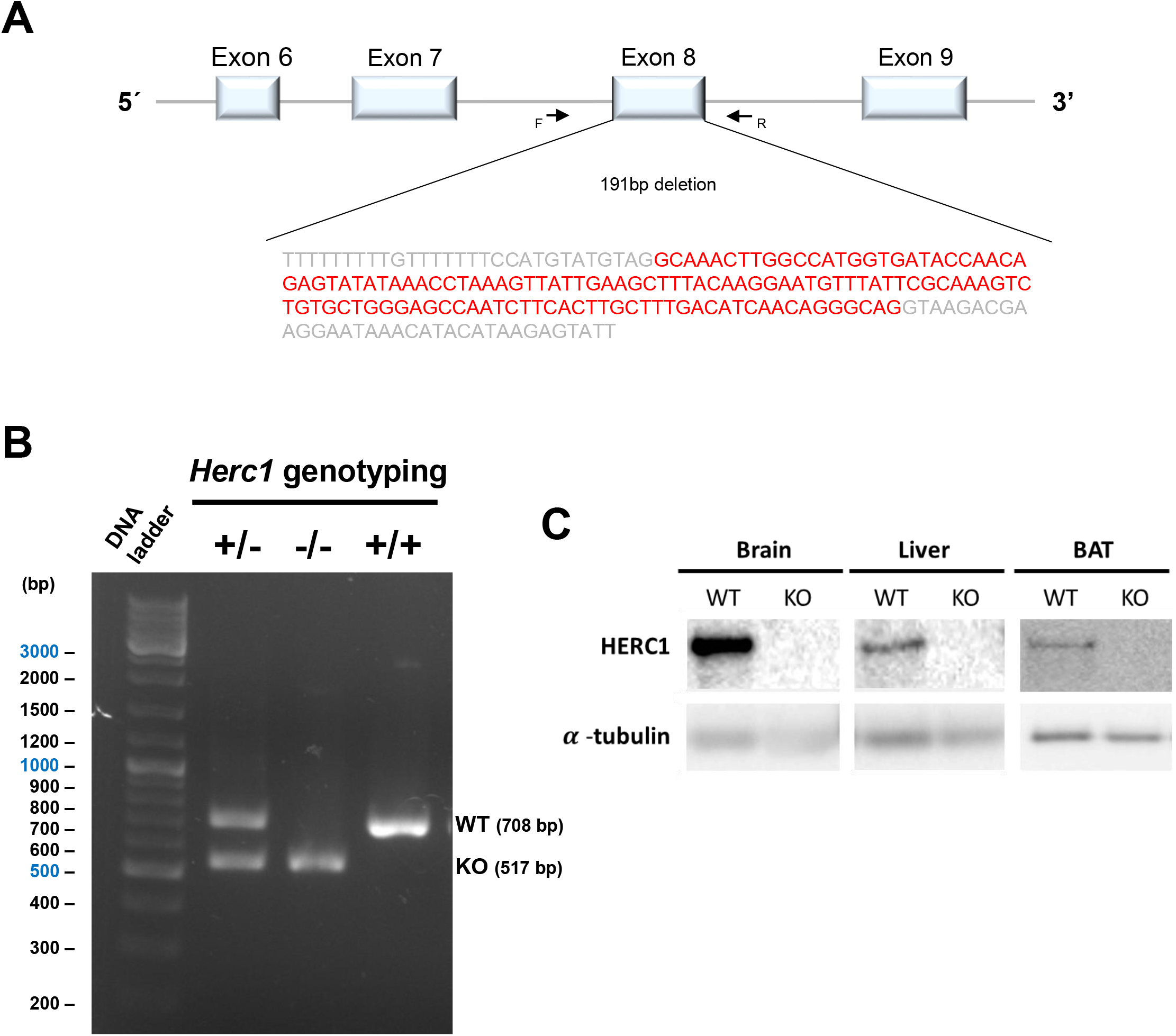
HERC1 loss-of-function mouse model. (A) Schematic representation of the targeted genomic region. A 191-bp deletion removes exon 8 of the *Herc1* gene on chromosome 9, generating the *Herc1*^em3(IMPC)Wtsi^ mutant allele. (B) Genotyping of wild-type (WT: +) and *Herc1* knockout (KO:-) variants by PCR. (C) HERC1 protein expression in tissues from WT and KO mice analysed by Western blot. HERC1 protein was undetectable in KO tissues, confirming loss of HERC1 expression. BAT: Brown adipose tissue. α-tubulin was used as a loading control.

**Supplementary Fig. 3.**
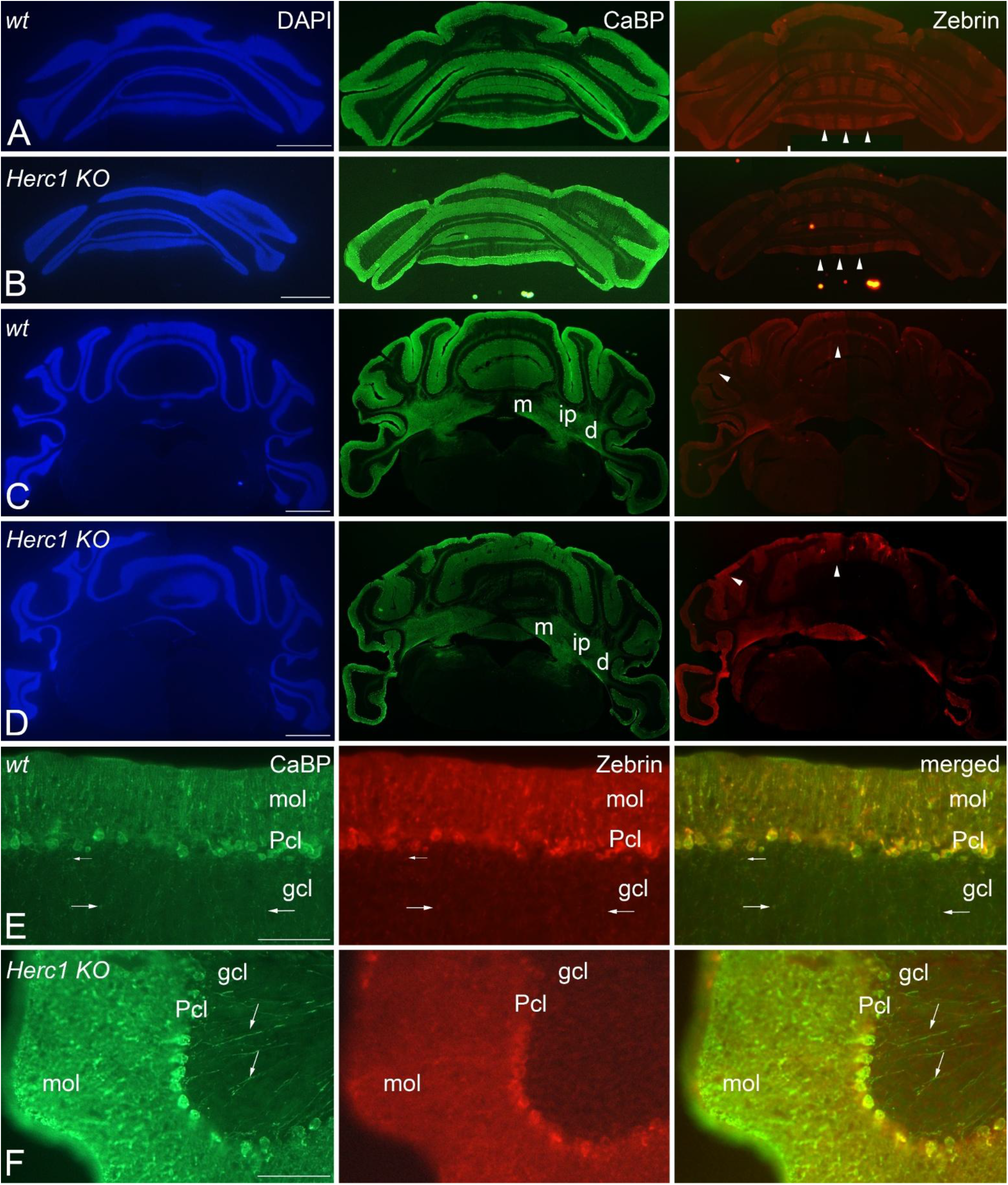
HERC1 deficiency does not compromise Purkinje cells viability or disrupt their monolayer parasagittal compartmentation. Montages of microphotographs of coronal sections through the cerebellum of control (**A**, **C** and **E**) and *Herc1 KO* (**B**, **D**, **F**) mice. The absence of HERC1 does not affect Purkinje cells survival and monolayer organization, as it shows their calbindin immunoreactivity (CaBP), which preserve their parasagittal bundles organization of Zebrin II immunoreactive Purkinje cells (arrowheads). The CaBP axonal plexus (**E**-**F**, small arrows) and the axons of the Purkinje cells (**E**-**F**, arrows) are devoid of the anomalous axonal torpedoes frequently found in HERC1 overproduction mutation (*see ref 21*). d, dentate nucleus; gcl, granule cell layer; ip, interposed nuclei; m, medial nucleus; mol, molecular layer; Pcl, Purkinje cell layer. Bars = 1mm (A-D), and 100 µm (E-F). As comparable results were observed in male and female mice, the data are presented collectively.

**Supplementary Fig. 4.**
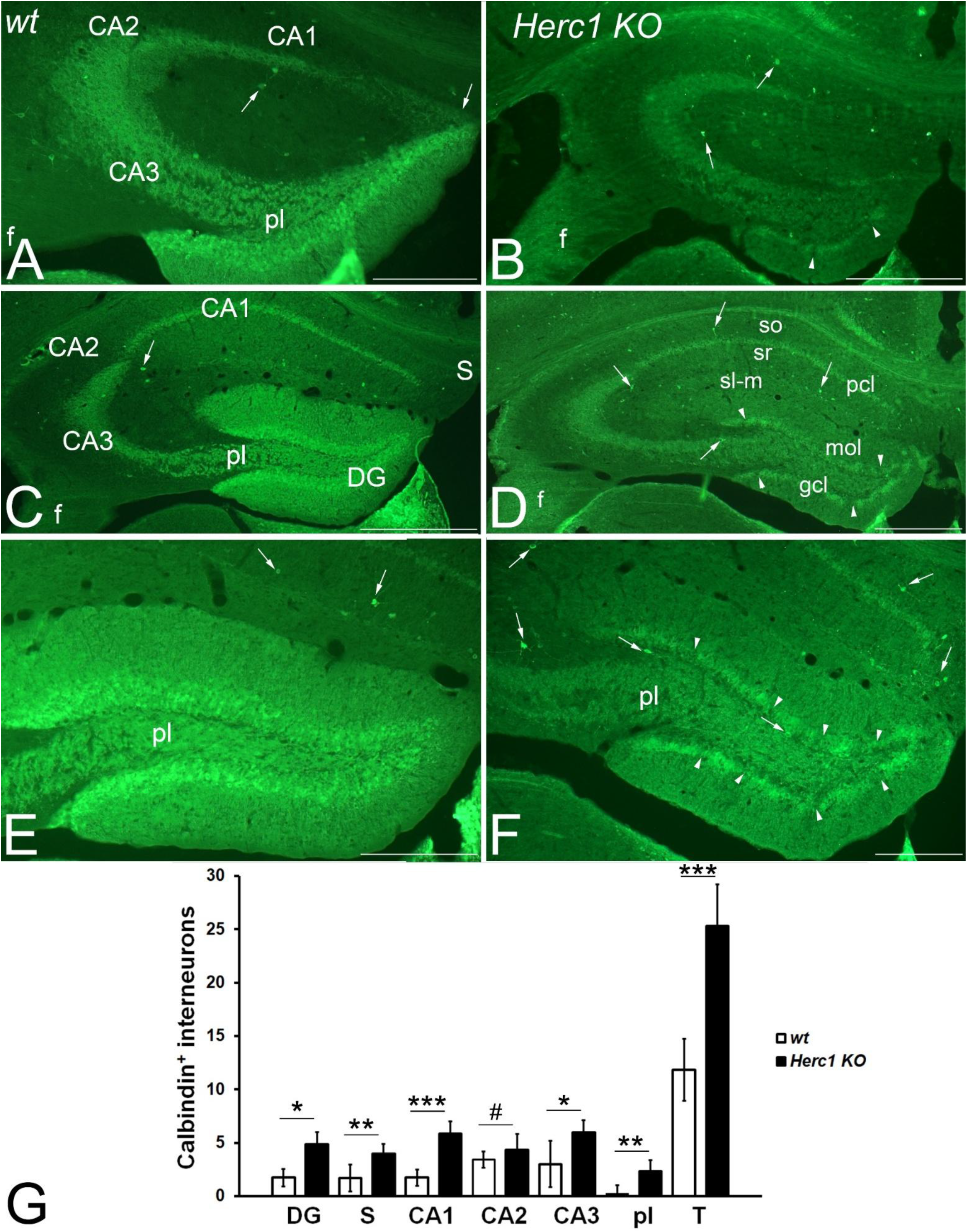
HERC1 deficiency increases calbindin immunoreactive interneurons of hippocampus. Coronal sections through the brain of control (**A, C, E**) and *Herc1 KO* (**B, D, F**) two-month-old mice. The granule cells layer (gcl) of the dentate gyrus (DG) appears thinner (**B, D, F**), than that of the hippocampus control (**A, C, E**), and displays patches devoid of calbindin immunoreactivity (**B, D, F**, arrowheads). A qualitative comparison of CaBP immunoreactive neurons shows that they are most abundant in the *KO* hippocampus (arrows). Quantitative analysis demonstrates that the *Herc1 KO* hippocampus has significantly more calbindin immunoreactive interneurons than the control (**G**; CA1, *p*= 0.000232; CA3, *p* = 0.010224; DG, *p* = 0.010738; S, *p* = 0.002976; pl, *p* = 0.002980) and only at the level of CA2 the difference is under the value of statistical significance (**G;** CA2, *p* = 0.225207). In present study any *p*-value less than 0.05 was considered significant, indicated as follows: \**p* < 0.05, \*\**p* < 0.01, and \*\*\**p* < 0.001. No significant values were indicated as follows: #*p* > 0.05. CA1, cornu ammonis 1; CA2, cornu ammonis 2; CA3, cornu ammonis 3; DG, dentate gyrus; S, subiculum; pcl, pyramidal cell layer; pl, polymorphic cell layer; sl-m, stratum lucidum-moleculare; so, stratum oriens; sr, stratum radiatum; T, total hippocampus. Bars = 500 µm (**A-D**) and 250 µm (**E-F**). No sex-differences were observed (*p* > 0.05), and the data are presented collectively.

**Supplementary Fig. 5.**
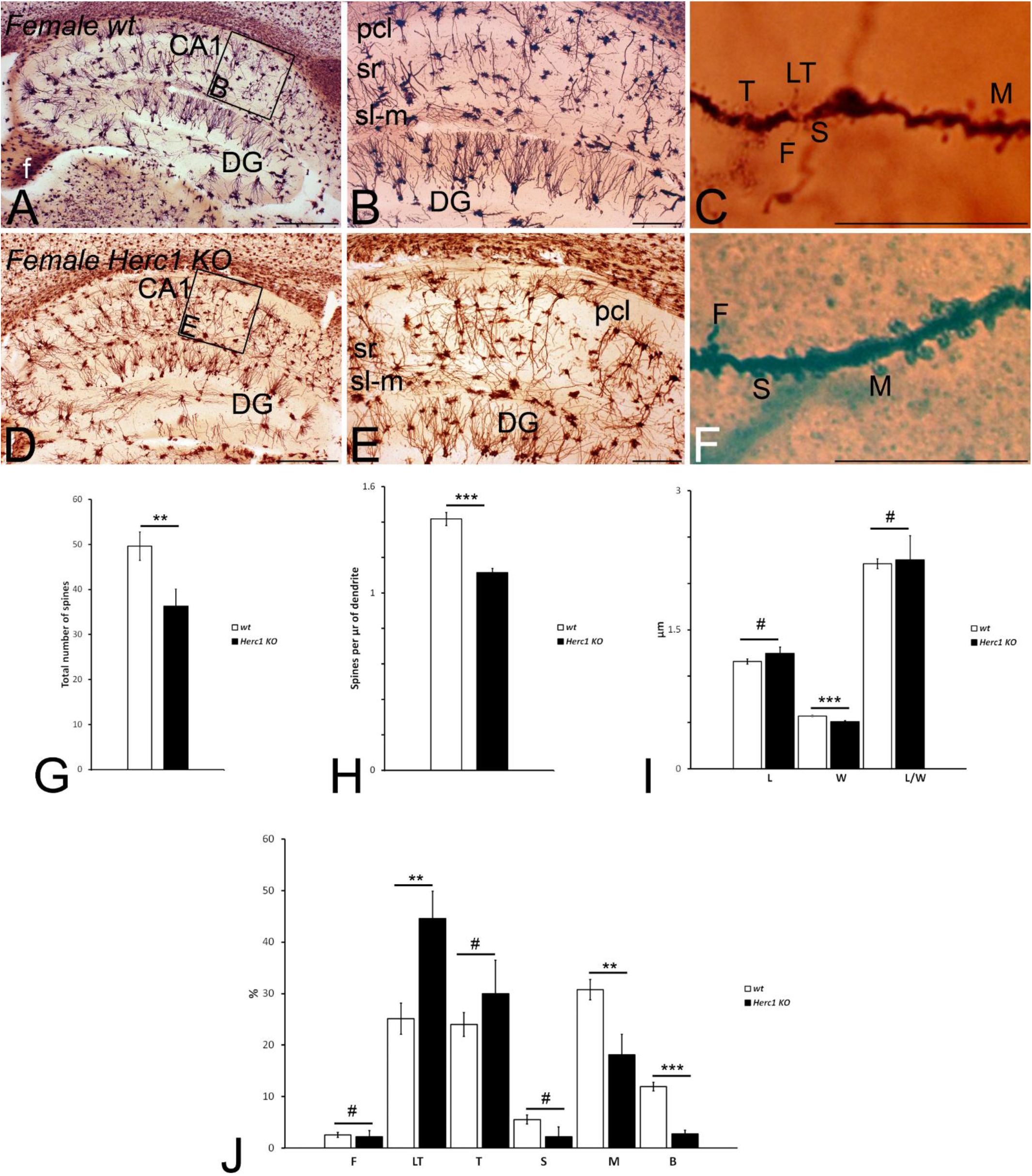
HERC1 deficiency alters dendritic spine morphology in hippocampal CA1 pyramidal neurons of female mice. Microphotographs of parasagittal sections through a Golgi-Cox impregnated hippocampus of control (**A**-**C**) and *Herc1 KO* (**D**-**F**) female mice. **C** and **F** Illustrate the fully filled dendritic segments used to count and categorize the spines. **G**–**J**, graphical representations of the number of spines counted (**G**, *p* = 0.0158894), the density of spines (**H**, *p* = 0.000013), and the spine differences in length (**I**, L, *p* = 0.260219), width (**I**, W, *p* = 0.000435), and length/width ratio (**I**, L/W Ratio, *p* = 0.873835) in the CA1 of wild-type and KO mice [**J**; *p* = 0.475506 (F), *p* = 0.330478 (LT), *p* = 0.434248 (T), *p* = 0.010673 (S), *p* = 0.003155 (M), *p* = 6.202806^E-06^ (B)]. The asterisks indicate significant differences in the Student’s t-test: \**p* < 0.05, \*\**p* < 0.01, and \*\*\**p* < 0.001. No significant values were indicated as follows: #*p* > 0.05. CA1, cornu ammonis 1; DG, dentate gyrus; f, fimbria; pcl, pyramidal cells layer; sl-m, stratum lacunosum-moleculare; sr, stratum radiatum. F, filopodia; LT, long thin spines; T, thin spines; M, mushroom spines; S, stubby spines; B, branched spines. Bars = 500 µm (A-B, D-E), and 50 µm (C, F).

**Supplementary Fig. 6.**
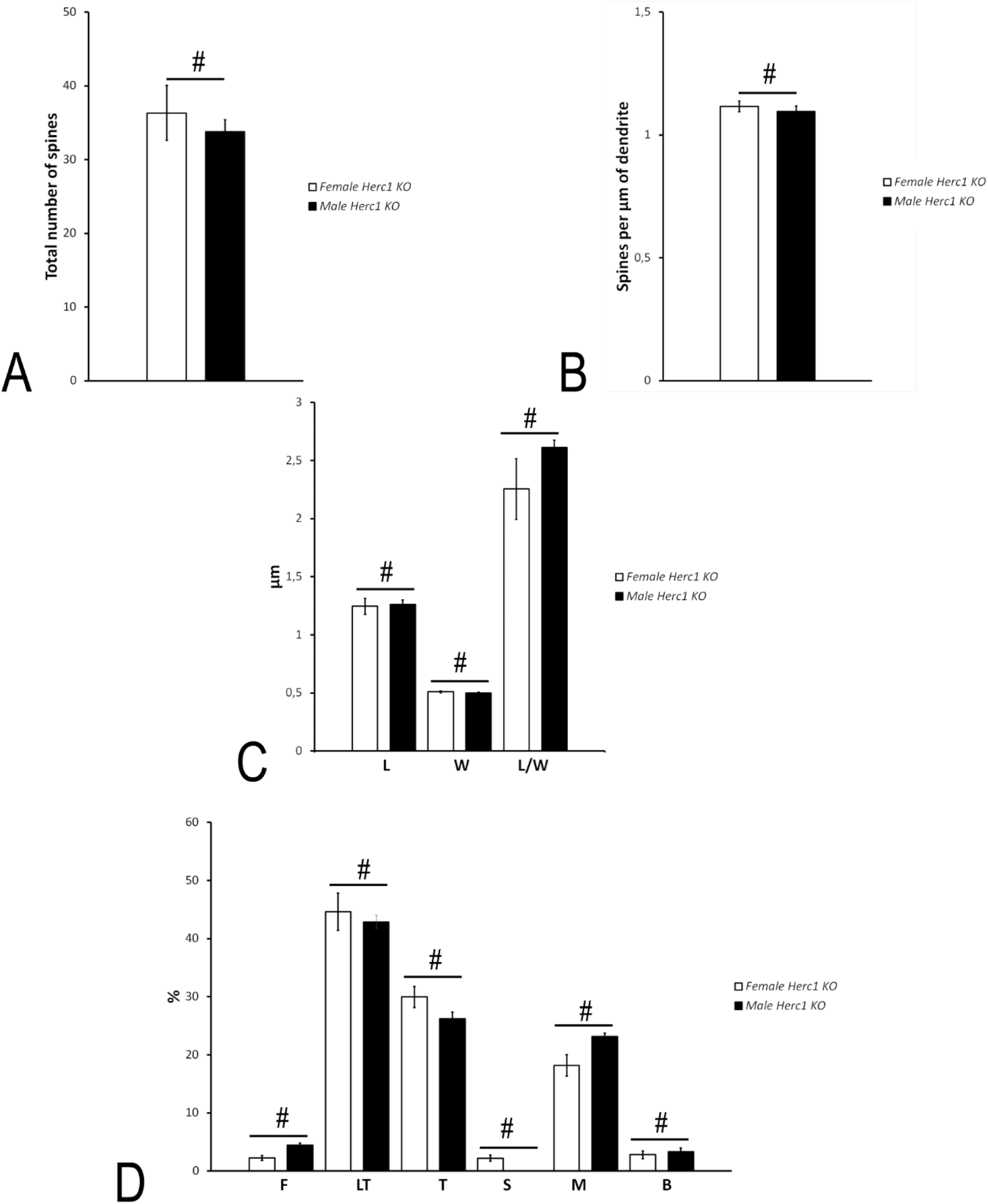
No sex-related differences in dendritic spine morphology of CA1 pyramidal neurons in *Herc1 KO* mice. Comparative analysis of the data shown in Fig. 3 (male mice) and Supplementary Fig. 4 (female mice) revealed no statistically significant differences in dendritic spine number, density, morphology, or subtype distribution between sexes. [**A**, *p* = 0.5424; **B**, *p* = 0.5294; **C**, *p* = 0.8625 (L), *p* = 0.41843 (W), *p* = 0.2190 (L/W); **D**, *p* = 0.1977 (F), *p* = 0.7642 (LT), *p* = 0.6088 (T), *p* = 0.2774 (S), *p* = 0.2691 (M), *p* = 0.6041 (B)]. L, length; W, width; L/W, length/width ratio; F, filopodia; LT, long thin spines; T, thin spines; M, mushroom spines; S, stubby spines; B, branched spines. # = *p* > 0.05.

**Supplementary Fig. 7.**
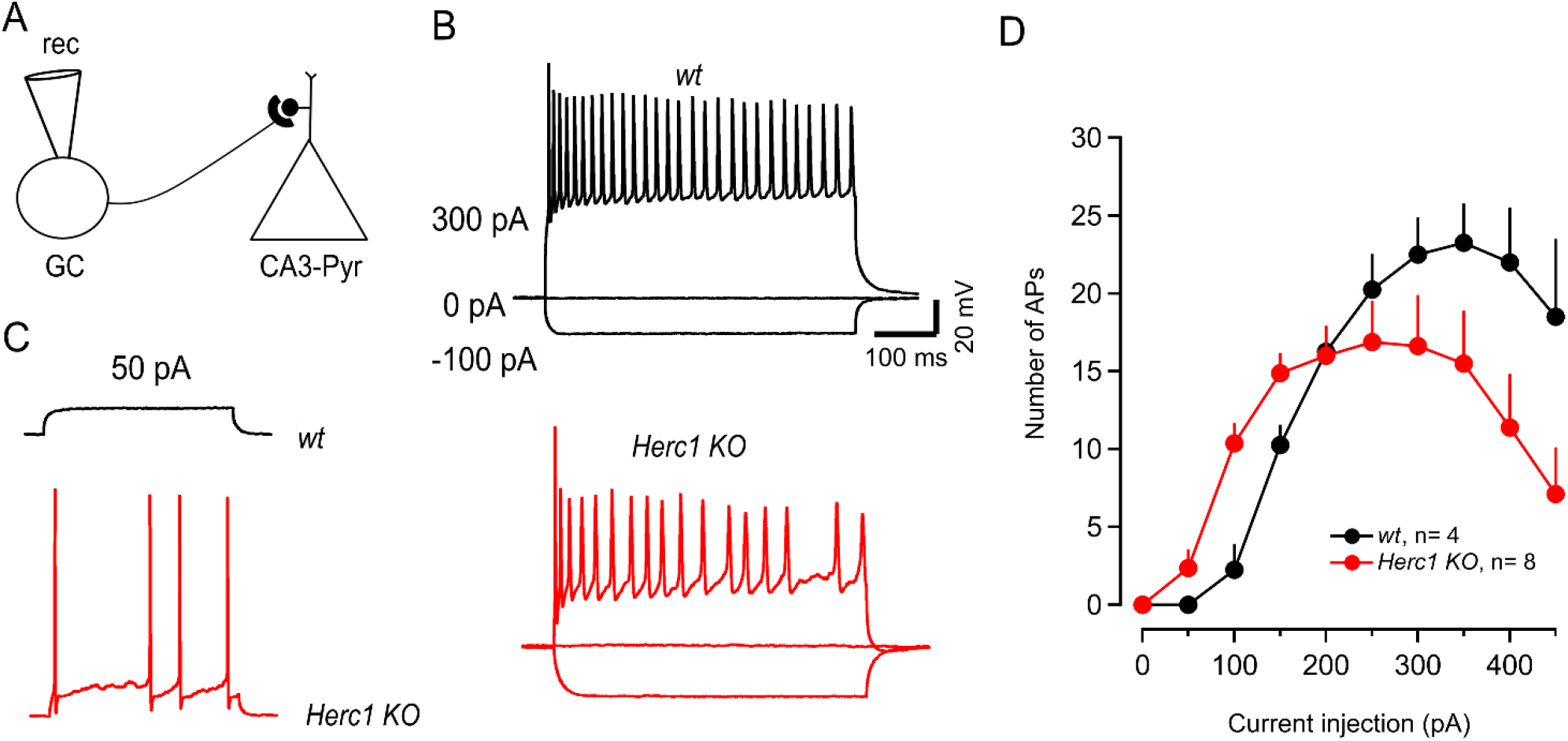
HERC1 deficiency impairs action potential firing in hippocampal granule cells. (A) Experimental electrophysiological configuration illustrating whole-cell current-clamp recording from a hippocampal granule cell GC to assess intrinsic excitability and action potential AP firing. GCs were subjected to 500 ms current-injection steps delivered in 50 pA increments, ranging from −250 to +450 pA. (B) Representative current-clamp recordings from a control (*wt*, top) and *Herc1 KO* (bottom) GC in response to −100, 0, and +300 pA current injections. Note the occurrence of AP failures in the *Herc1 KO* GC. (C) Representative current-clamp recordings illustrating differences in membrane excitability between *wt* and *Herc1 KO* GCs in response to a +50 pA current injection. (D) Mean number of APs elicited during 500 ms current-injection steps delivered in 50-pA increments from 0 to +450 pA in *wt* and *Herc1 KO* GCs.

**Supplementary Fig. 8.**
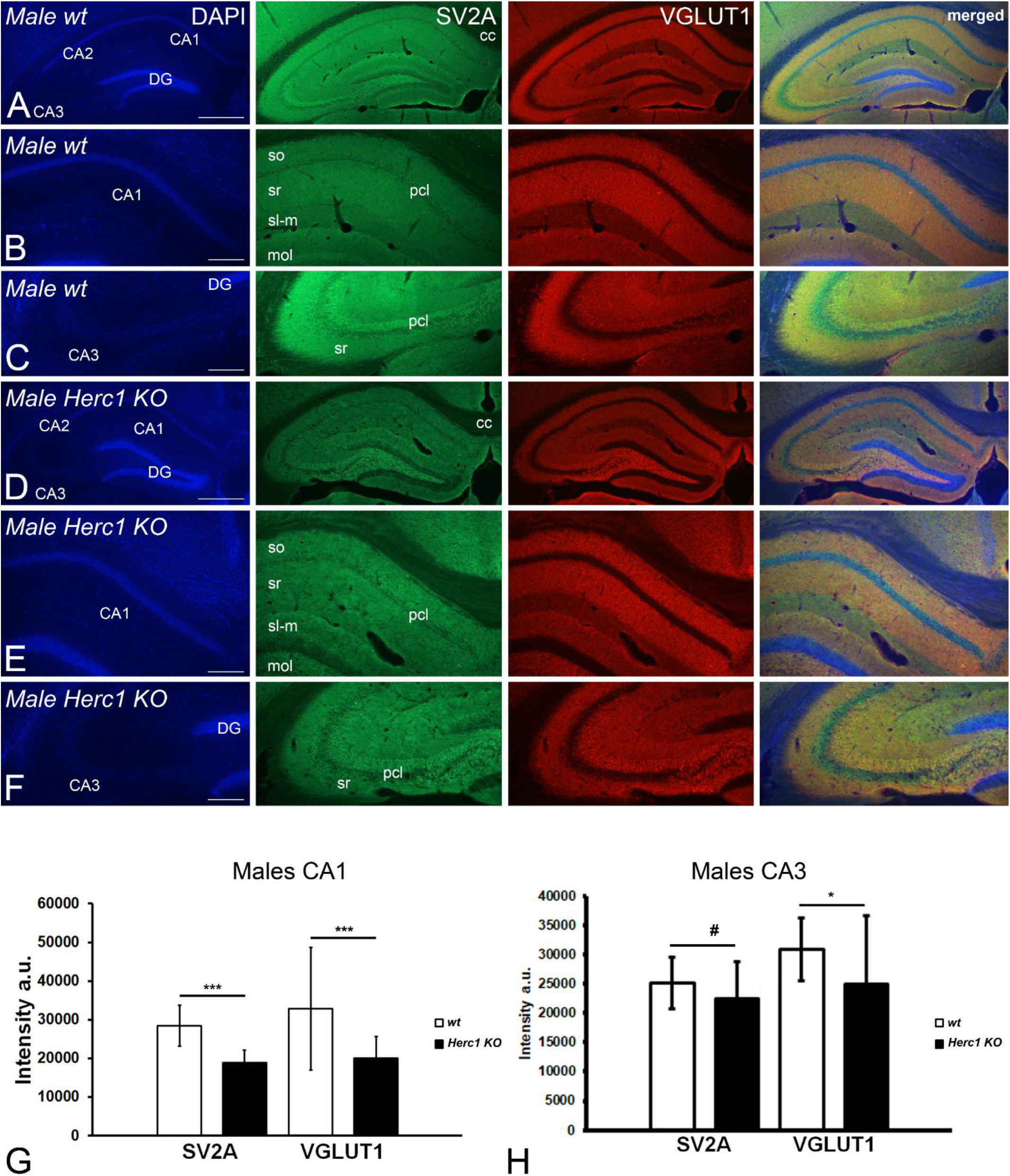
HERC1 deficiency decreases the presynaptic proteins SV2 and VGLT1 expression in male hippocampus. Coronal sections through the hippocampus of two-month-old male control (A-C) and *Herc1 KO* (D-F) mice. Confocal laser images illustrate the expression of synaptic vesicle protein 2 (SV2A) and the vesicular glutamate transporter 1 (VGLUT1) in the entire hippocampus (A and D), CA1 (B and E) and CA3 (C and F) regions. Note as the intensity of the immunoreactivity in control hippocampus is higher than in *Herc1 KO* one. Quantitative analyses demonstrate that presynaptic proteins expression of control mice is strongest related to *Herc1 KO* ones [G, *p* = 9.41E-11 (SV2A) and *p* = 0.000116 (VGLUT1); and H, *p* = 0.057267 (SV2A) and *p* = 0.012091 (VGLUT1)]. The asterisks indicate significant differences in the Student’s t-test: \**p* < 0.05, \*\**p* < 0.01, and \*\*\**p* < 0.001. CA1. #*p* >0.05. CA1, cornu ammonis 1; CA2, cornu ammonis 2; CA3, cornu ammonis 3; DG, dentate gyrus; cc, corpus callosum; mol, molecular layer of the dentate gyrus; pcl, pyramidal cells layer; sl-m, stratum lacunosum-moleculare; so, stratum oriens; sr, stratum radiatum. Bars = 500 µm (A and D), and 200 µm (B-C, E-F).

**Supplementary Fig. 9.**
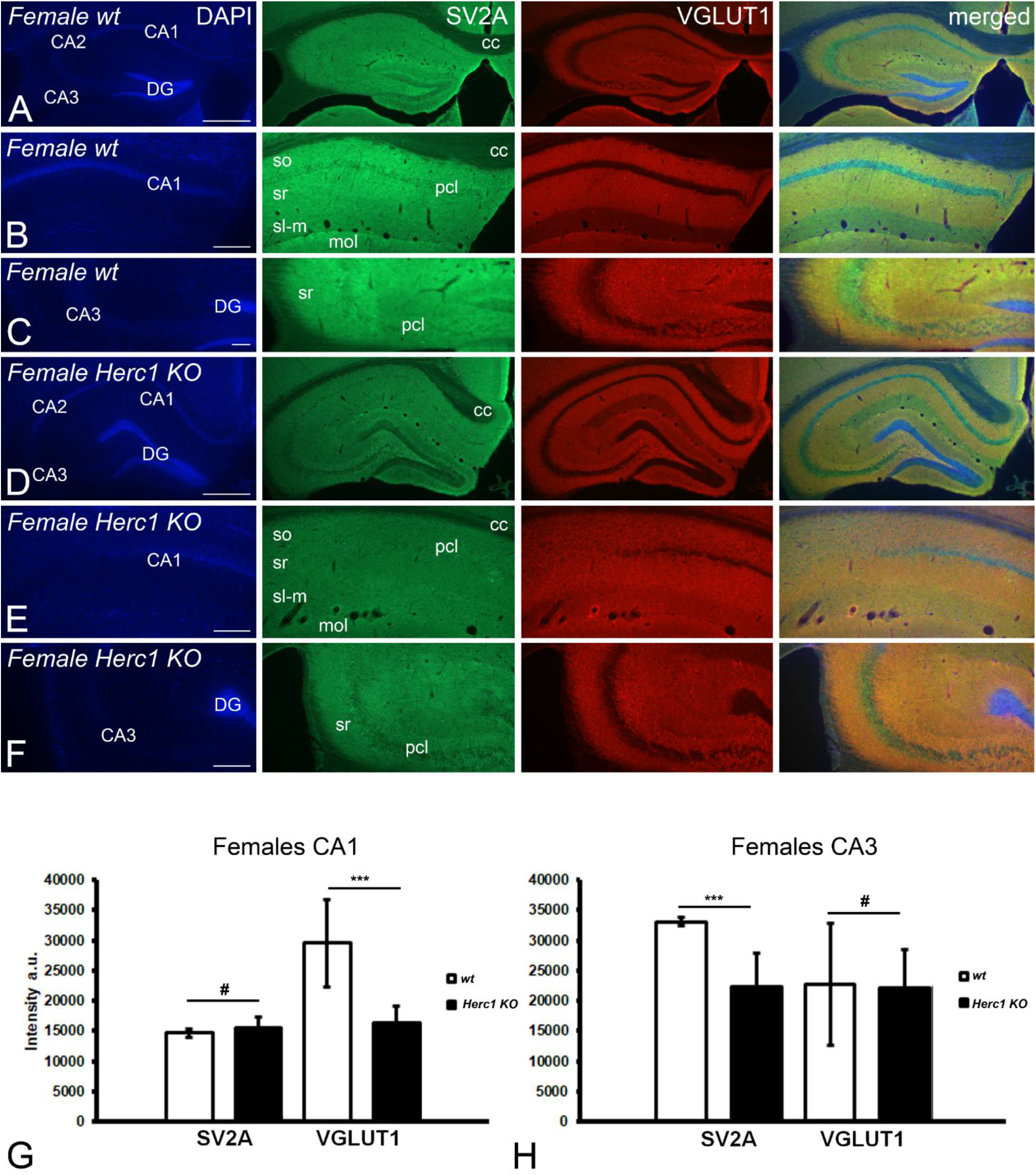
HERC1 deficiency induces a discrete decrease in the presynaptic proteins SV2 and VGLT1 expression in female hippocampus. Coronal sections through the hippocampus of two-month-old female control (A-C) and *Herc1 KO* (D-F) mice. Confocal laser images illustrate the expression of synaptic vesicle protein 2 (SV2A) and the vesicular glutamate transporter 1 (VGLUT1) in the entire hippocampus (A and D) and in CA1 (B and E) CA3 (C and F) areas. Note as the intensity of the immunoreactivity in Herc1 KO hippocampus seems to be slightly lesser than in control one. Quantitative analyses only show statistically significant differences in the SV2 expression at CA3 (G, *p* = 4.48E-07) while these differences stand out for VGLUT1 at CA1 area (H, p = 1.122981E-12). The asterisks indicate significant differences in the Student’s t-test: \**p* < 0.05, \*\**p* < 0.01, and \*\*\**p* < 0.001. #*p* >0.05. CA1, cornu ammonis 1; CA2, cornu ammonis 2; CA3, cornu ammonis 3; DG, dentate gyrus; cc, corpus callosum; mol, molecular layer of the dentate gyrus; pcl, pyramidal cells layer; sl-m, stratum lacunosum-moleculare; so, stratum oriens; sr, stratum radiatum. Bars = 500 µm (A and D), and 200 µm (B-C, E-F).

**Supplementary Fig. 10.**
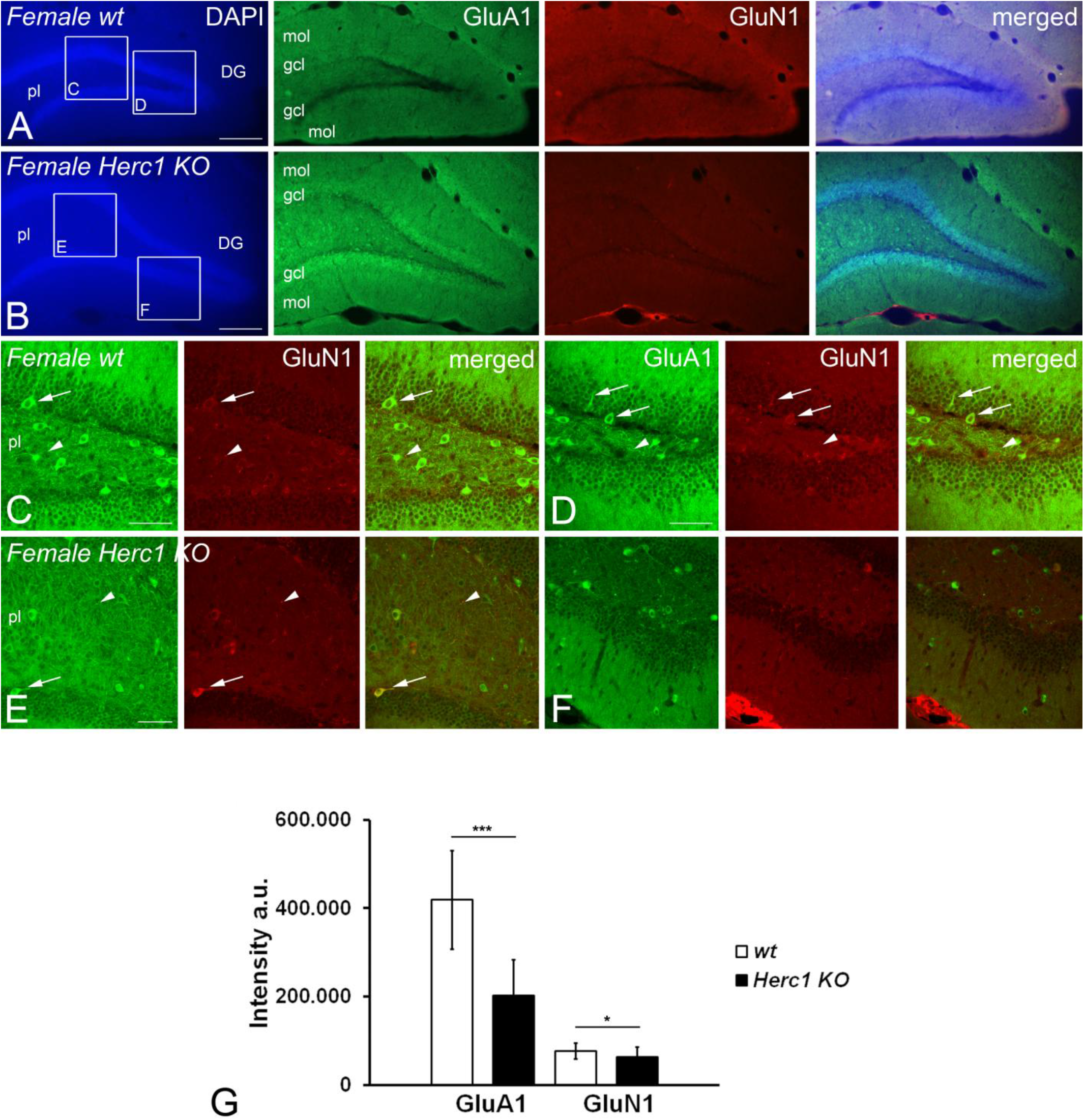
HERC1 deficiency decreases the neurons expressing postsynaptic proteins in female dentate gyrus. Coronal sections through the hippocampus of two-month-old female control (A, C-D) and *Herc1 KO* (B, E-F) mice. Confocal laser images illustrate granule cells neurons of the dentate gyrus (C, D, E) and neurons of the polymorphic layer of the dentate gyrus (C-F) which express the subunit GluA1 of the AMPA ionotropic receptor (GluA1, green) and the GluN1 subtype of NMDA receptor (GluN1, red) or both (arrows) (C-D, E-F). Double labelled presynaptic boutons can be also observed (arrowheads) in this region (C, D, E). Note as immunoreactive neurons are more abundant in control (C-D) than in *Herc1 KO* (E-F) polymorphic layer (pl). The quantitative decrease of both postsynaptic proteins in Herc1 KO female dentate gyrus is statistically significant regarding to the control (G, *p* = 5.83^E-05^ for GluA1, and *p* = 0.0408 for GluN1). The asterisks indicate significant differences in the Student’s t-test: \**p* < 0.05, \*\**p* < 0.01, and \*\*\**p* < 0.001. DG, dentate gyrus; gcl, granule cells layer; mol, molecular layer of the dentate gyrus; pl, polymorphic layer of the dentate gyrus. Bars = 500 µm (A and D), and 200 µm (B-C, E-F).

**Supplementary Fig. 11.**
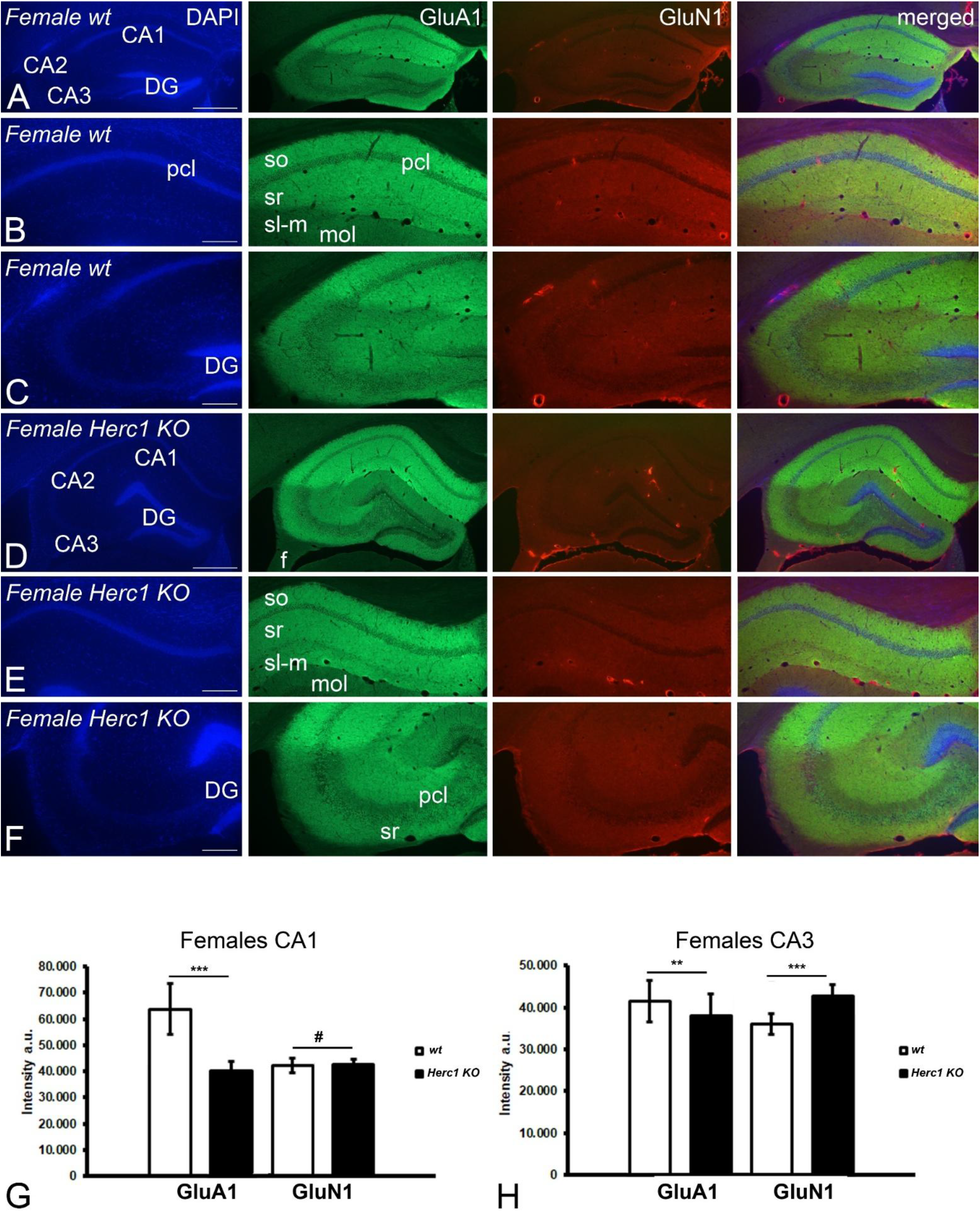
HERC1 deficiency disturbs the postsynaptic proteins GluA1 and GluN1 expression in female hippocampus. Coronal sections through the hippocampus of two-month-old female control (A-C) and *Herc1 KO* (D-F) mice. Confocal laser images illustrate the expression of the subunit GluA1 of the AMPA ionotropic receptor (GluA1) and the GluN1 subtype of NMDA receptor (GluN1) in the entire hippocampus (A and D), CA1 (B and E) and CA3 (C and F) regions. Note as the intensity of the immunoreactivity in control and *Herc1 KO* hippocampus seems to be qualitatively similar. Furthermore, the clearcut of GluA1 expression at the border between CA1 and CA2 hippocampal regions of the control brains (A), also appears in *Herc1 KO* ones (D). Quantitative analyses show that GluA1 expressions are stronger in control hippocampus (G, *p* = 5.54388E-15; H, *p* = 0.00696) than in *Herc1 KO* one. On the contrary, GluN1 immunoreactivity is most intense in *Herc1 KO* CA3 (H, *p* = 1.19605E-15) while no varies regarding the control in the CA1 area (G, *p* = 0.6126). The asterisks indicate significant differences in the Student’s t-test: \**p* < 0.05, \*\**p* < 0.01, and \*\*\**p* < 0.001. #*p* >0.05. CA1, cornu ammonis 1; CA2, cornu ammonis 2; CA3, cornu ammonis 3; DG, dentate gyrus; mol, molecular layer of the dentate gyrus; pcl, pyramidal cells layer; sl-m, stratum lacunosum-moleculare; so, stratum oriens; sr, stratum radiatum. Bars = 500 µm (A and D), and 200 µm (B-C, E-F).

**Supplementary Fig. 12.**
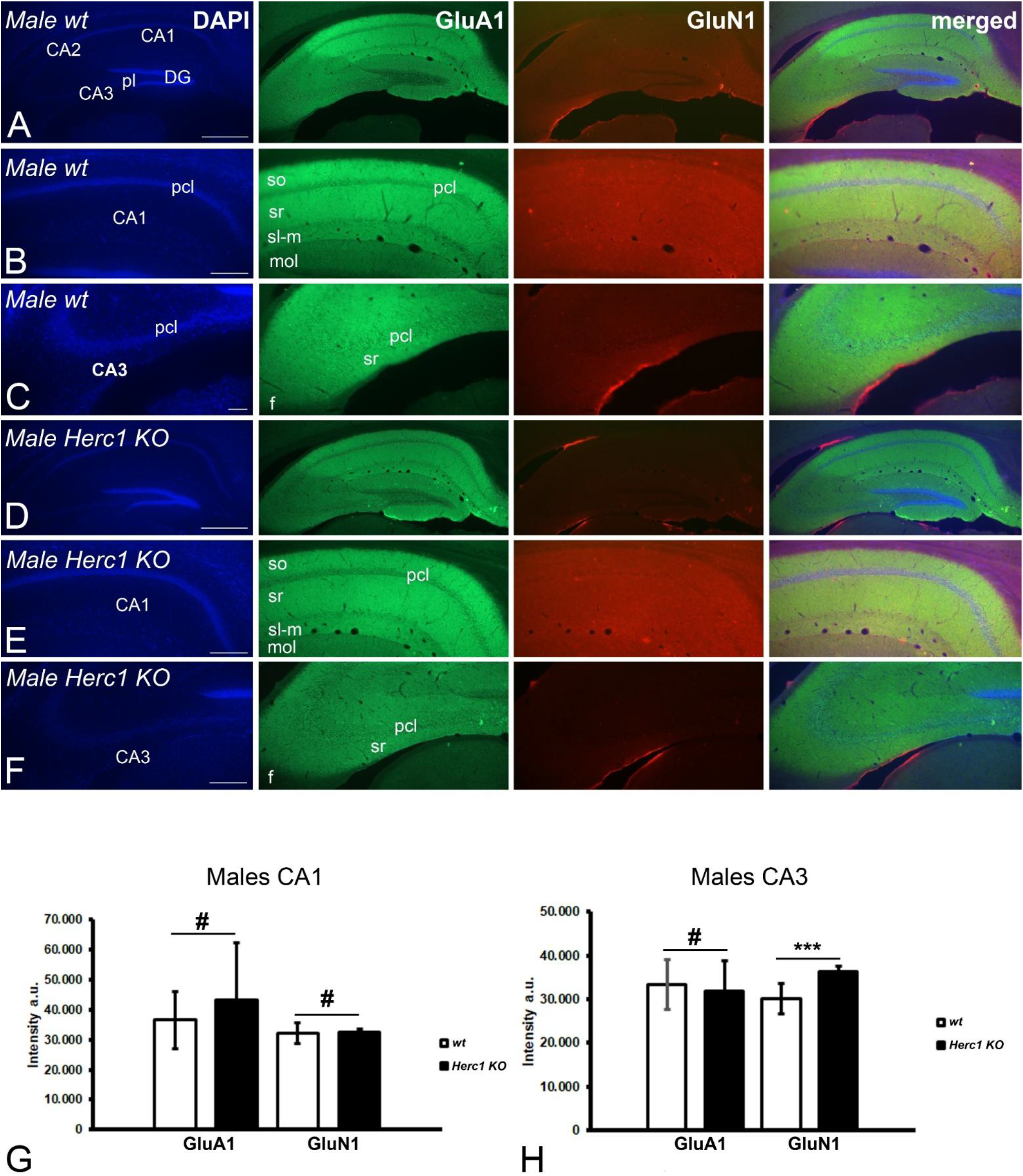
HERC1 deficiency induces few changes in the expression of postynaptic proteins GluA1 and GluN1 in male hippocampus. Coronal sections through the hippocampus of two-month-old male control (A-C) and *Herc1 KO* (D-F) mice. Confocal laser images illustrate the expression of the subunit GluA1 of the AMPA ionotropic receptor (GluA1) and the GluN1 subtype of NMDA receptor (GluN1) in the entire hippocampus (A and D), the CA1 (B and E) and CA3 (C and F) regions. Note as the intensity of the immunoreactivity in control and *Herc1 KO* hippocampus seems to be qualitatively similar. Furthermore, the clearcut of GluA1 expression at the border between CA1 and CA2 observed in female hippocampus is also present in of the control (A) and *Herc1 KO* male brains (D). Quantitative analyses show that GluN1 expressions are stronger in *Herc1 KO* CA3 (H, p = 4.73561E-11) than in control one, while values of the expression of GluN1 in CA1 and GluA1 in both CA1 and CA3 are under the level of significance [G, p = 0.0906 (GluA1); p = 0.438116763 (GluN1); H, p = 0.364947162 (GluA1)]. The asterisks indicate significant differences in the Student’s t-test: \**p* < 0.05, \*\**p* < 0.01, and \*\*\**p* < 0.001. #*p* >0.05. CA1, cornu ammonis 1; CA2, cornu ammonis 2; CA3, cornu ammonis 3; DG, dentate gyrus; f, fimbria; mol, molecular layer of the dentate gyrus; pcl, pyramidal cells layer; sl-m, stratum lacunosum-moleculare; so, stratum oriens; sr, stratum radiatum. Bars = 500 µm (A and D), and 200 µm (B-C, E-F).

**Supplementary Fig. 13.**
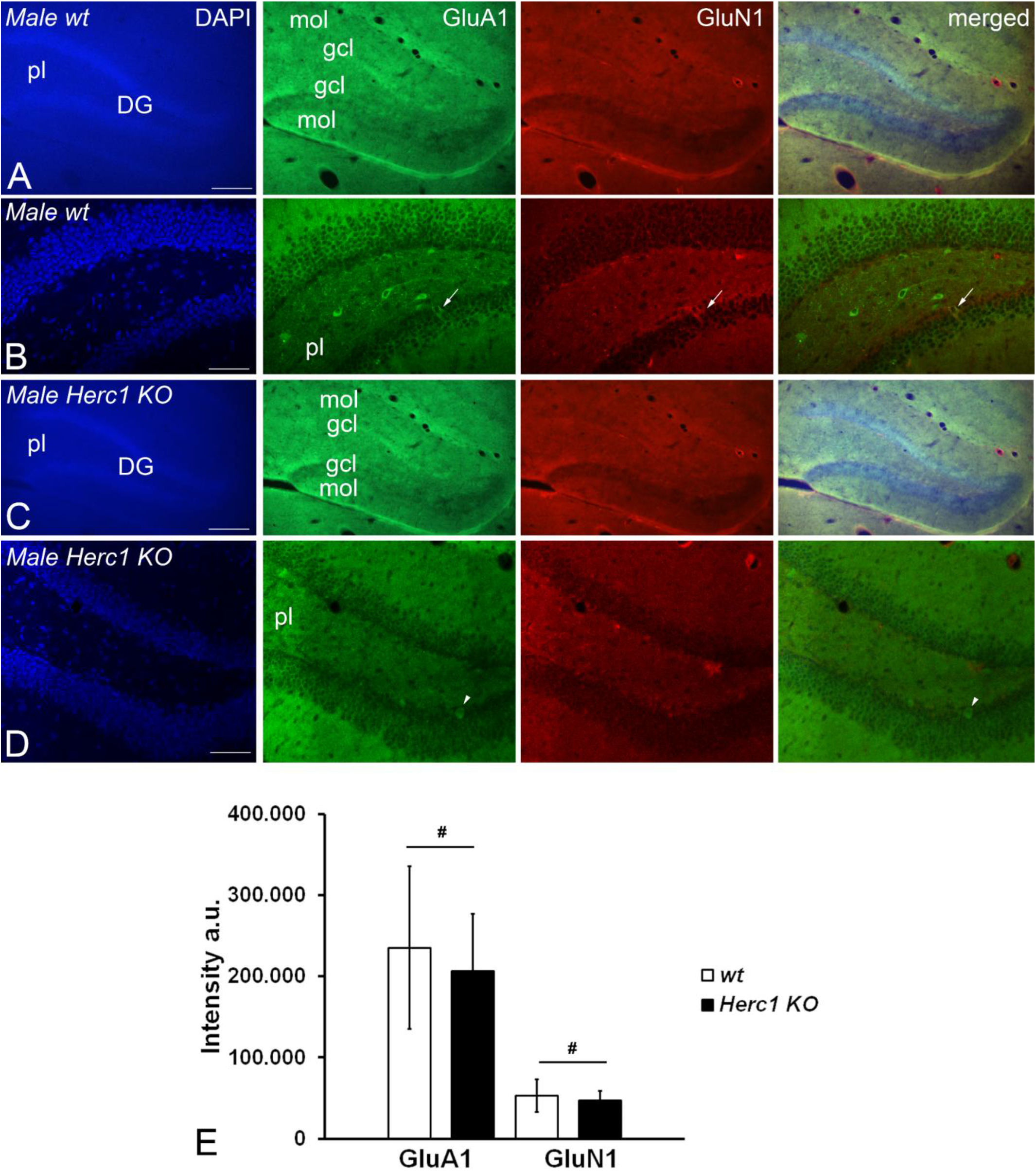
HERC1 deficiency does not influence the neurons expressing postsynaptic proteins in male dentate gyrus. Coronal sections through the hippocampus of two-month-old male control (A, C) and *Herc1 KO* (B, D) mice. Confocal laser images illustrate granule cells neurons of the dentate gyrus (B, D) which express the subunit GluA1 of the AMPA ionotropic receptor (GluA1, B, arrow; D, arrowhead) and the GluN1 subtype of NMDA receptor (GluN1, B, arrow) or both (B, arrow). As in the female mice the immunoreactive neurons are more abundant in control (C-D) than in *Herc1 KO* (E-F) polymorphic layer (pl); however, it should be highlighted that qualitatively they are less numerous than in the female dentate gyrus (see Supplementary Figure 10). The quantitative analysis does not find statistically significant differences in the expression of both postsynaptic proteins (E, *p* = 0.4877 (GluA1), and *p* = 0.4845 (GluN1)). # = *p* >0.05. DG, dentate gyrus; gcl, granule cells layer; mol, molecular layer of the dentate gyrus; pl, polymorphic layer of the dentate gyrus. Bars = 500 µm (A and D), and 200 µm (B-C, E-F).

**Supplementary Fig. 14.**
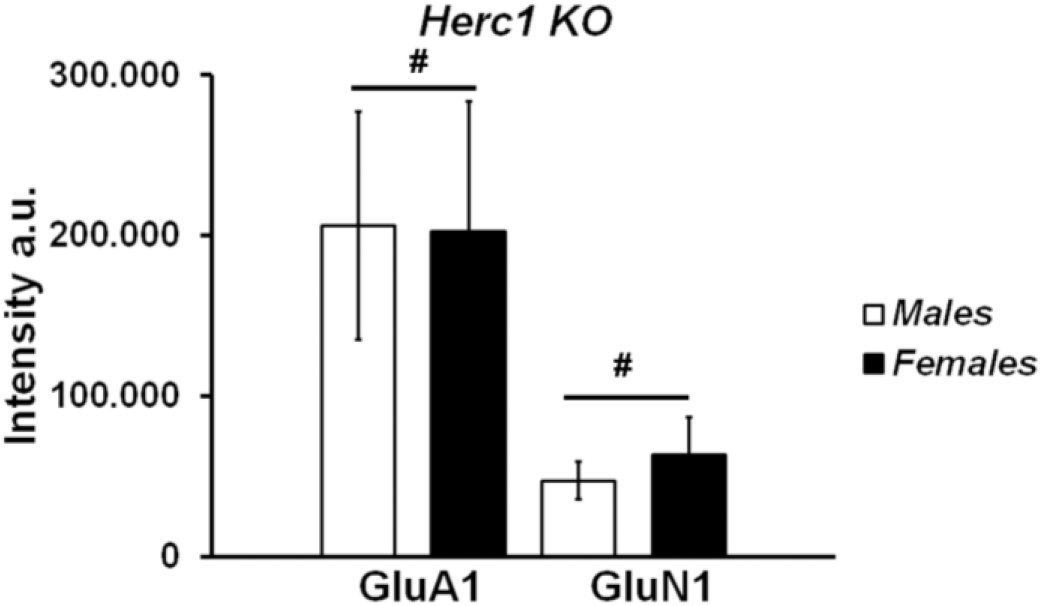
No sex-related differences in the expression of GluA1 and GluN1 in the dentate gyrus. Graphic illustrates the comparison between the levels of expression of the subunit GluA1 of the AMPA ionotropic receptor (GluA1) and the GluN1 subtype of NMDA receptor (GluN1). Although apparently *Herc1KO* female shows less immunoreactivity to both glutamate receptor markers, the difference is under the level of significance. (#, *p* = 0.9212 for GluA1 and *p* = 0.07213 for GluN1).

## References

1. Reshadmanesh A, Dehdahsi S, Ahangari F, et al. First Case of Macrocephaly, Dysmorphic Facies, and Psychomotor Retardation Harboring Co-inherited Variants in HERC1 and PMP22 Genes from Iran: Two Novel Variants. Arch Iran Med. 2024;27(12):700–706. doi:10.34172/aim.31593

2. Aggarwal S, Bhowmik A Das, Ramprasad VL, Murugan S, Dalal A. A splice site mutation in HERC1 leads to syndromic intellectual disability with macrocephaly and facial dysmorphism: Further delineation of the phenotypic spectrum. Am J Med Genet A. 2016;170(7):1868–1873. doi:10.1002/AJMG.A.37654

3. Utine GE, Taşkıran EZ, Koşukcu C, et al. HERC1 mutations in idiopathic intellectual disability. Eur J Med Genet. 2017;60(5):279–283. doi:10.1016/J.EJMG.2017.03.007

4. Nguyen LS, Schneider T, Rio M, et al. A nonsense variant in HERC1 is associated with intellectual disability, megalencephaly, thick corpus callosum and cerebellar atrophy. European Journal of Human Genetics. 2016;24(3):455–458. doi:10.1038/EJHG.2015.140

5. Schwarz JM, Pedrazza L, Stenzel W, Rosa JL, Schuelke M, Straussberg R. A new homozygous HERC1 gain-of-function variant in MDFPMR syndrome leads to mTORC1 hyperactivation and reduced autophagy during cell catabolism. Mol Genet Metab. 2020;131(1-2):126–134. doi:10.1016/J.YMGME.2020.08.008

6. Pérez-Villegas EM, Ruiz R, Bachiller S, Ventura F, Armengol JA, Rosa JL. The HERC proteins and the nervous system. Semin Cell Dev Biol. Published online November 27, 2021. doi:10.1016/j.semcdb.2021.11.017

7. Ortega-Recalde O, Beltrán OI, Gálvez JM, et al. Biallelic HERC1 mutations in a syndromic form of overgrowth and intellectual disability. Clin Genet. 2015;88(4):e1–e3. doi:10.1111/CGE.12634

8. Mashimo T, Hadjebi O, Amair-Pinedo F, et al. Progressive Purkinje cell degeneration in tambaleante mutant mice is a consequence of a missense mutation in HERC1 E3 ubiquitin ligase. PLoS Genet. 2009;5(12):e1000784. doi:10.1371/journal.pgen.1000784

9. Pérez-Villegas EM, Negrete-Díaz J V., Porras-García ME, et al. Mutation of the HERC 1 Ubiquitin Ligase Impairs Associative Learning in the Lateral Amygdala. Mol Neurobiol. 2018;55(2):1157–1168. doi:10.1007/S12035-016-0371-8

10. Pérez-Villegas EM, Pérez-Rodríguez M, Negrete-Díaz J V., et al. HERC1 Ubiquitin Ligase Is Required for Hippocampal Learning and Memory. Front Neuroanat. 2020;14. doi:10.3389/FNANA.2020.592797

11. Pedrazza L, Martinez-Martinez A, Sánchez-de-Diego C, et al. HERC1 deficiency causes osteopenia through transcriptional program dysregulation during bone remodeling. Cell Death Dis. 2023;14(1). doi:10.1038/s41419-023-05549-x

12. Cubillos-Rojas M, Schneider T, Hadjebi O, et al. The HERC2 ubiquitin ligase is essential for embryonic development and regulates motor coordination. Oncotarget. 2016;7(35):56083–56106. doi:10.18632/oncotarget.11270

13. Võikar V, Stanford SC. The Open Field Test. Neuromethods. 2023;190:9–29. doi:10.1007/978-1-0716-2748-8_2

14. Guyenet SJ, Furrer SA, Damian VM, et al. A Simple Composite Phenotype Scoring System for Evaluating Mouse Models of Cerebellar Ataxia. J Vis Exp. 2010;(39):e1787. doi:10.3791/1787

15. Deacon RMJ. Measuring motor coordination in mice. J Vis Exp. 2013;(75). doi:10.3791/2609

16. Can A, Dao DT, Terrillion CE, Piantadosi SC, Bhat S, Gould TD. The tail suspension test. J Vis Exp. 2012;(59). doi:10.3791/3769

17. Hawkes R, Herrup K. Aldolase C/zebrin II and the regionalization of the cerebellum. J Mol Neurosci. 1995;6(3):147–158. doi:10.1007/BF02736761

18. Clarke PGH. An unbiased correction factor for cell counts in histological sections. J Neurosci Methods. 1993;49(1-2):133–140. doi:10.1016/0165-0270(93)90117-A

19. Risher WC, Ustunkaya T, Alvarado JS, Eroglu C. Rapid Golgi analysis method for efficient and unbiased classification of dendritic spines. PLoS One. 2014;9(9). doi:10.1371/JOURNAL.PONE.0107591

20. Small SA, Schobel SA, Buxton RB, Witter MP, Barnes CA. A pathophysiological framework of hippocampal dysfunction in ageing and disease. Nat Rev Neurosci. 2011;12(10):585–601. doi:10.1038/NRN3085

21. Koziol LF, Budding D, Andreasen N, et al. Consensus paper: the cerebellum’s role in movement and cognition. Cerebellum. 2014;13(1):151–177. doi:10.1007/S12311-013-0511-X

22. Elena Porras-García M, Ruiz R, Pérez-Villegas EM, Armengol JÁ. Motor learning of mice lacking cerebellar Purkinje cells. Front Neuroanat. 2013;(MARCH). doi:10.3389/FNANA.2013.00004

23. Pelkey KA, Chittajallu R, Craig MT, Tricoire L, Wester JC, McBain CJ. Hippocampal gabaergic inhibitory interneurons. Physiol Rev. 2017;97(4):1619–1747. doi:10.1152/physrev.00007.2017

24. Tzilivaki A, Tukker JJ, Maier N, Poirazi P, Sammons RP, Schmitz D. Hippocampal GABAergic interneurons and memory. Neuron. 2023;111(20):3154–3175. doi:10.1016/j.neuron.2023.06.016

25. Witter MP, Amaral DG. Hippocampal Formation. The Rat Nervous System. Published online January 1, 2004:635–704. doi:10.1016/B978-012547638-6/50022-5

26. Jinno S, Kosaka T. Cellular architecture of the mouse hippocampus: A quantitative aspect of chemically defined GABAergic neurons with stereology. Neurosci Res. 2006;56(3):229–245. doi:10.1016/j.neures.2006.07.007

27. Ramón y Cajal S. Textura del sustema nervioso del hombre y de los vertebrados: estudios sobre el plan estructural y compsicion histologica de los centros nerviosos adicionados de consideraciones fisiologicas fundadas en los neuvos descubrimientos: Ramón y Cajal, Santi…. Re-imp. Gráficas Vidal Leuca, SL, Alicante, SPAIN, 1992. ISBN: 84-604-4778-2. 1904. Accessed October 30, 2026. https://archive.org/details/b21270855

28. Freund TF, Buzsáki G. Interneurons of the Hippocampus. Hippocampus. 1996;6(4):347–470. doi:10.1002/(sici)1098-1063(1996)6:4<347::aid-hipo1>3.0.co;2-i

29. Kempermann G, Song H, Gage FH. Neurogenesis in the Adult Hippocampus. Cold Spring Harb Perspect Biol. 2015;7(9):a018812. doi:10.1101/CSHPERSPECT.A018812

30. Liu SH, Wang J, Zhu DY, Fu YP, Lukowiak K, Lu YM. Generation of functional inhibitory neurons in the adult rat hippocampus. J Neurosci. 2003;23(3):732–736. doi:10.1523/JNEUROSCI.23-03-00732.2003

31. Hogg JA, Cousin MA. Control of Synaptotagmin-1 Trafficking by SV2A-Mechanism and Consequences for Presynaptic Function and Dysfunction. J Neurochem. 2025;169(1). doi:10.1111/JNC.16308

32. Du X, Li J, Li M, et al. Research progress on the role of type I vesicular glutamate transporter (VGLUT1) in nervous system diseases. Cell Biosci. 2020;10(1). doi:10.1186/S13578-020-00393-4

33. Diering GH, Huganir RL. The AMPA Receptor Code of Synaptic Plasticity. Neuron. 2018;100(2):314–329. doi:10.1016/j.neuron.2018.10.018

34. Carles A, Freyssin A, Perin-Dureau F, Rubinstenn G, Maurice T. Targeting N-Methyl-d-Aspartate Receptors in Neurodegenerative Diseases. Int J Mol Sci. 2024;25(7). doi:10.3390/IJMS25073733

35. Kannan M, Bayam E, Wagner C, et al. WD40-repeat 47, a microtubule-associated protein, is essential for brain development and autophagy. Proceedings of the National Academy of Sciences. 2017;114(44):E9308–E9317. doi:10.1073/PNAS.1713625114

36. Doludda B, Barde W, D’Egidio F, et al. Adult neurogenesis: New neurons, new opportunities. Cell Stem Cell. 2026;33(3):382–392. doi:10.1016/j.stem.2026.01.014

37. Wyllie DJA, Bowie D. Ionotropic glutamate receptors: structure, function and dysfunction. J Physiol. 2022;600(2):175–179. doi:10.1113/JP282389

38. Iyer L, Johnson K, Collier S, Koretsky AP, Petrus E. Post-Critical Period Transcriptional and Physiological Adaptations of Primary Sensory Cortex after Sensory Loss. J Neurosci. 2026;46(7). doi:10.1523/JNEUROSCI.1519-25.2025

39. Acsády L, Kamondi A, Sík A, Freund T, Buzsáki G. GABAergic cells are the major postsynaptic targets of mossy fibers in the rat hippocampus. J Neurosci. 1998;18(9):3386–3403. doi:10.1523/JNEUROSCI.18-09-03386.1998

40. Smith SS, Waterhouse BD, Chapin JK, Woodward DJ. Progesterone alters GABA and glutamate responsiveness: a possible mechanism for its anxiolytic action. Brain Res. 1987;400(2):353–359. doi:10.1016/0006-8993(87)90634-2

41. Pettorossi VE, Di Mauro M, Scarduzio M, et al. Modulatory role of androgenic and estrogenic neurosteroids in determining the direction of synaptic plasticity in the CA1 hippocampal region of male rats. Physiol Rep. 2013;1(7):e00185. doi:10.1002/phy2.185

42. Sakata K, Tokue A, Kawai N. Altered synaptic transmission in the hippocampus of the castrated male mouse is reversed by testosterone replacement. Journal of Urology. 2000;163(4):1333–1338. doi:10.1016/S0022-5347(05)67773-7

43. Kuwahara N, Nicholson K, Isaacs L, MacLusky NJ. Androgen Effects on Neural Plasticity. Androgens. 2021;2(1):216–230. doi:10.1089/ANDRO.2021.0022

44. Hatanaka Y, Hojo Y, Mukai H, et al. Rapid increase of spines by dihydrotestosterone and testosterone in hippocampal neurons: Dependence on synaptic androgen receptor and kinase networks. Brain Res. 2015;1621:121–132. doi:10.1016/j.brainres.2014.12.011

45. Hu W, Macdonald ML, Elswick DE, Sweet RA. The glutamate hypothesis of schizophrenia: evidence from human brain tissue studies. Ann N Y Acad Sci. 2015;1338(1):38–57. doi:10.1111/NYAS.12547

46. Singh T, Poterba T, Curtis D, et al. Rare coding variants in ten genes confer substantial risk for schizophrenia. Nature 2022 604:7906. 2022;604(7906):509-516. doi:10.1038/s41586-022-04556-w

47. Li W, Ghose S, Gleason K, et al. Synaptic proteins in the hippocampus indicative of increased neuronal activity in CA3 in schizophrenia. Am J Psychiatry. 2015;172(4):373–382. doi:10.1176/APPI.AJP.2014.14010123

